# T cell-specific P2RX7 favors lung parenchymal CD4^+^ T cell accumulation in response to severe lung infections

**DOI:** 10.1101/2022.09.19.508603

**Authors:** Igor Santiago-Carvalho, Gislane Almeida-Santos, Bruna Gois Macedo, Caio Cesar Barbosa-Bomfim, Fabricio Moreira Almeida, Marcos Vinícios Pinheiro Cione, Trupti Vardam-Kaur, Mia Masuda, Sarah Van Dijk, Bruno Marcel Melo, Rogério Silva do Nascimento, Rebeka da Conceição Souza, Alba Lucínia Peixoto-Rangel, Robson Coutinho-Silva, Mario H. Hirata, José Carlos Alves-Filho, José Maria Álvarez, Elena Lassounskaia, Henrique Borges da Silva, Maria Regina D’Império-Lima

**Author notes:** Correspondence (H.B.D.S) and (M.R.D.L). These authors have contributed equally to this work (Henrique Borges da Silva, Maria Regina D’Império-Lima).

## Abstract

CD4^+^ T cells are key components of the immune response during lung infections and can mediate protection against tuberculosis (TB) or influenza. However, CD4^+^ T cells can also promote lung pathology during these infections, making it unclear how these cells control such discrepant effects. Using mouse models of hypervirulent TB and influenza, we observed that exaggerated accumulation of parenchymal CD4^+^ T cells promotes lung damage. Low numbers of lung CD4^+^ T cells, in contrast, are sufficient to protect against hypervirulent TB. In both situations, lung CD4^+^ T cell accumulation is mediated by CD4^+^ T cell-specific expression of the extracellular ATP (eATP) receptor P2RX7. P2RX7 upregulation in lung CD4^+^ T cells promotes expression of the chemokine receptor CXCR3 and favors *in situ* proliferation. Our findings suggest that direct sensing of lung eATP by CD4^+^ T cells is critical to induce tissue CD4^+^ T cell accumulation and pathology during lung infections.

**GRAPHICAL ABSTRACT:** 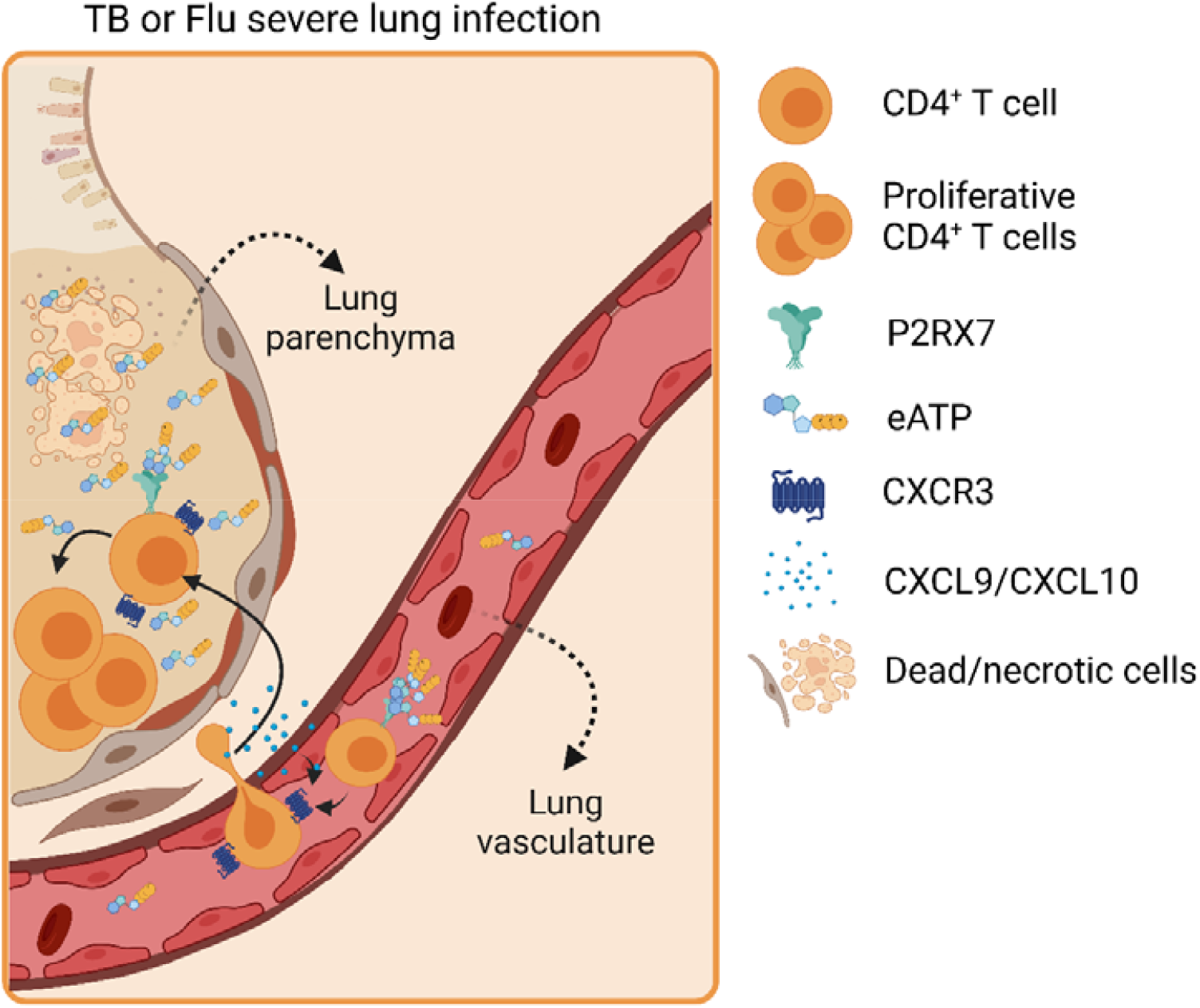

## INTRODUCTION

The magnitude of immunopathology caused by infectious diseases is a consequence of the intensity and duration of the immune response (Medzhitov et al., 2012). The strength of the immune response is also determined by the tissue sensitivity to damage and its regenerative capacity. In the lung, for example, there is a fine line between an effective or harmful immune response (Medzhitov et al., 2012; Ravimohan et al., 2018). A dysregulated inflammatory response in the lung can cause severe tissue damage and compromise respiratory physiology (Klomp et al., 2021; Ravimohan et al., 2018). CD4^+^ T cells are key players in the immune response of many lung diseases (Brown et al., 2004; McDermott and Klein, 2018; Peng et al., 2020; Sakai et al., 2014). That includes infections such as tuberculosis (TB), one of the world’s deadliest bacterial diseases (WHO, 2021), and influenza, which still is a major inducer of patient hospitalizations and deaths across the world (Paget et al., 2019). During these infections, accumulation of distinct CD4^+^ T cell subsets mediates protection or worsening of the disease (Moguche et al., 2017; Riberdy et al., 2000). Pathogen-specific effector CD4^+^ T cells can either remain in the circulation and populate the lung intravascular regions or infiltrate the lung parenchyma (Anderson et al., 2014; Sakai et al., 2014). The cell-intrinsic molecular basis underlying these homing choices has been subject of recent studies, showing for example the importance of differential chemokine receptor expression between these two subsets. In response to virulent H37Rv *Mycobacterium tuberculosis* (*Mtb*), CD4^+^ T cells expressing the fractalkine receptor CX3CR1 tend to stay within the vasculature and produce high levels of IFN-γ, while cells expressing the CXC Motif Chemokine Receptor 3 (CXCR3) tend to establish parenchymal residency and are more effective in containing infection despite lower IFN-γ production in a per-cell-basis (Sakai et al., 2014; Sallin et al., 2017). CXCR3 also promotes lung CD4^+^ T cell residency and subsequent protection in response to influenza (Shane et al., 2020). The balance between protective immune responses promoted by CD4^+^ T cells, which are by definition pleiotropic, and their unintended effects in the lung parenchyma defines whether parenchymal CD4^+^ T cells are protective or harmful (Ehlers, 1999).

The influence of lung microenvironmental factors in the generation of parenchymal CD4^+^ T cell populations is less understood. However, Influenza and *Mtb* infect distinct parenchymal cells (epithelial cells and macrophages, respectively), and consequently induce different local immune responses and changes in the lung microenvironment (O’Garra et al., 2013; Thomas et al., 2006). *Mtb* often induces the generation of complex immune/parenchymal cell structures called granulomas (Nunes-Alves et al., 2014; Saunders et al., 1999), while flu leads to diffuse accumulation of immune cells around infected areas (Damjanovic et al., 2011; Peiris et al., 2010). In both infections, however, inflammatory responses are triggered by pathogen load, local cell death and immune cell infiltration. An important inflammation primer is the release of damage-associated molecular patterns (DAMPs), which can contribute to the dysregulation of lung microenvironment and immune responses (Gong et al., 2020; Patel, 2018). Extracellular ATP (eATP) is one of the most abundant DAMPs released in these processes (Idzko et al., 2007; Kono and Rock, 2008). High levels of eATP are recognized by the low-affinity purinergic receptor P2RX7 which is mostly expressed on immune cells, including CD4^+^ T cells (Schenk et al., 2008; Surprenant et al., 1996). Our group has previously demonstrated that expression of P2RX7 in hematopoietic cells contributes to the generation of severe pulmonary TB (Amaral et al., 2014; Bomfim et al., 2017). Furthermore, pharmacological blockade of P2RX7 signaling prevents the generation of severe TB (Santiago-Carvalho et al., 2021). The biomedical importance of P2RX7 as a promoter of lung disease is further suggested by a previous report in influenza-infected global P2RX7-deficient mice (Leyva-Grado et al., 2017). Whether P2RX7 serves as a microenvironmental sensor for lung CD4^+^ T cell accumulation, and whether P2RX7-expressing CD4^+^ T cells are the main promoter of lung damage rather than other cell types is not understood. Of note, we have found that, in correlation with protection from disease, transient P2RX7 blockade led to reduced numbers of CD69^+^ lung-parenchymal CD4^+^ T cells (Santiago-Carvalho et al., 2021). Moreover, P2RX7 is known to play an important role in the establishment of tissue-resident memory (T_RM_) CD8^+^ T cells in response to systemic viruses (Borges da Silva et al., 2018; Borges da Silva et al., 2020).

Here, we report that cell-intrinsic P2RX7 promotes the lung parenchymal accumulation of effector CD4^+^ T cells in response to TB and influenza, directly promoting disease exacerbation. P2RX7-expressing CD4^+^ T cells increase CXCR3 expression and can access the lung parenchyma. In the lung parenchyma, P2RX7 promotes the proliferation of effector CD4^+^ T cells, further increasing their numbers in the tissue. Exaggerated (but not low) numbers of P2RX7^+^ parenchymal CD4^+^ T cells lead to disease promoting lung inflammation. Overall, our results suggest that sensing of lung-derived eATP is crucial to induce parenchymal CD4^+^ T cell establishment and exemplifies how the lung microenvironment shapes tissue-localized CD4^+^ T cell responses.

## RESULTS

### Parenchymal CD4^+^ T cells aggravate pulmonary TB caused by hypervirulent mycobacteria

We first infected C57BL/6 (WT) mice intratracheally with the hypervirulent mycobacterial *Mycobacterium bovis* (*Mbv*) MP287 strain (Amaral et al., 2014; Santiago-Carvalho et al., 2021). At day 28 p.i. MP287 infection induced severe lung damage, associated with extensive inflammatory infiltrates, intragranulomatous necrotic lesions, and areas of alveolitis with bronchial obstruction (**Figure 1A**). Alveoli were filled with inflammatory components, including monocytes, neutrophils, lymphocytes and fibrin. Areas of perifocal exudative reaction (hypersensitivity reaction) surrounding TB foci are marked by black arrows. It consists of edema and interstitial lymphocytic inflammation. We then assessed the intravascular and parenchymal CD4^+^ T cell infiltrate in the lung using intravenous anti-CD45 labeling (Anderson et al., 2014). In this context, at 28 days p.i., we noted a greater number of parenchymal (CD45i.v.^-^) than intravascular (CD45i.v.^+^) CD44^+^CD4^+^ T cells in the infected lung (**Figure 1B**). Phenotypic analysis of antigen-experienced (CD44^+^) CD4^+^ T cells revealed that KLRG1 and CX3CR1 molecules were mostly expressed on intravascular CD4^+^ T cells, while CD69 and CXCR3 were primarily expressed on parenchymal CD4^+^ T cells (**Figure S1A**), like what was previously described during *Mtb* H37Rv infection (Sakai et al., 2014).

**Figure 1.**
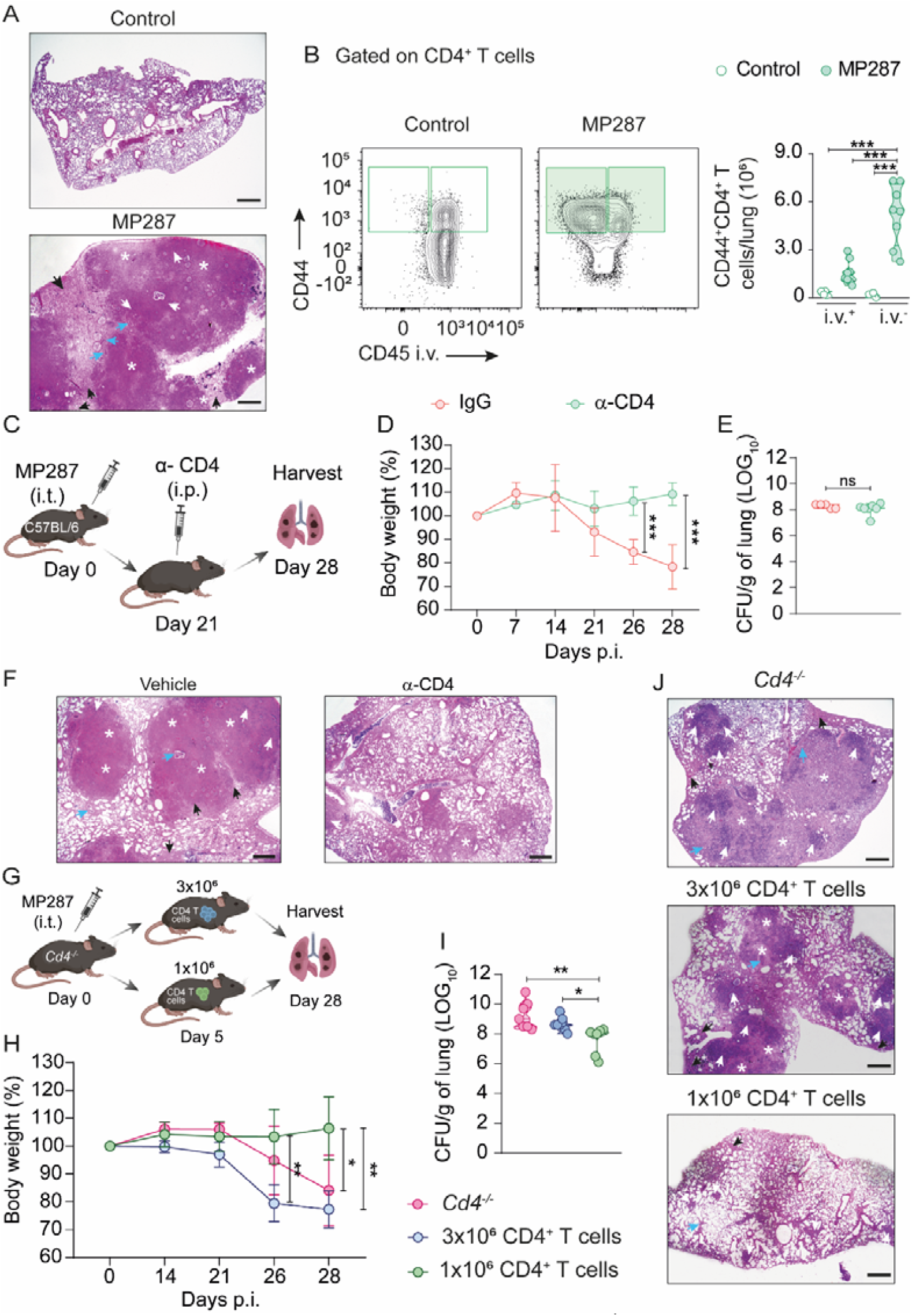
Excessive accumulation of parenchymal CD4^+^ T cells induces lung severe disease in response to hypervirulent mycobacteria. (**A-J**) C57BL/6 (WT) or CD4-KO mice (*Cd4*^*-/-*^) were infected with ∼100 CFU of the *Mbv*-MP287 strain. Non-infected mice were used as controls. At 28 days p.i., mice were injected i.v. with anti-CD45 fluorescent antibodies 3 minutes before lung harvest. (**A**) Representative lung sections of WT mice stained with hematoxylin-eosin (HE) (bar scales = 500 μM). White asterisks indicate areas of necrosis, white arrows indicate area of alveolitis, blue arrows indicate bronchial obstruction and black arrows indicate perifocal exudative reaction. (**B**) Left: flow cytometry plots of CD44 and CD45i.v. expression in CD4^+^ T cells of WT mice. Right: violin graphs of the numbers of CD45i.v.^+^ and CD45i.v.^-^ CD44^+^CD4^+^ T cells per lung. (**C**) Schematic illustration of CD4^+^ T cell depletion protocol (Created with BioRender.com). Infected WT mice were treated with control IgG or α-CD4 antibody. (**D**) Percentages of body weights in relation to day 0 of mice described in C. (**E**) CFU numbers per lung of mice described in C. (**F**) Representative lung sections stained with HE of mice described in C (bar scales = 500 μM). (**G**) Schematic illustration of adoptive transfer protocol (Created with BioRender.com). Infected *Cd4*^-/-^ mice were transferred or not with 1×10^6^ or 3×10^6^ CD4^+^ T cells. (**H**) Percentages of body weights in relation to day 0 of mice described in G. (**I**) CFU numbers per lung of mice described in G. (**J**) Representative lung sections stained with HE of mice described in G (bar scales = 500 μm). Data are from 2–3 independent experiments; n = 3-5 per experimental group per experiment. Graphical data is shown as means ± SD. p values from One-way ANOVA (Tukey post-tests) (**B, H** and **I**) or unpaired T-tests (**D-E**) are indicated by ^*^, p <0.05; ^**^, p<0.01; ^***^, p <0.001.

The contribution of massive infiltration of parenchymal CD44^+^CD4^+^ T cells to the worsening of TB was assessed by depleting CD4^+^ T cells with a high dose of α-CD4 antibody on day 21 p.i. (**Figures 1C and S1B**), when disease onset occurs in MP287-infected mice (Bomfim et al., 2017). CD4^+^ T cell depletion prevented weight loss induced by MP287 (**Figure 1D**). Despite this, depletion of CD4^+^ T cells did not change pulmonary CFUs (**Figure 1E**). Lung weights and total cell numbers were reduced in CD4^+^ T cell-depleted mice (**Figures S1C-D**). The size of the upper right lobe of the lung and the presence of multiple nodules were also decreased by CD4^+^ T cell depletion (**Figure S1E**), as well as the areas of pneumonia, alveolitis, bronchial obstruction and necrosis (**Figure 1F**).

To investigate whether lung parenchymal CD4^+^ T cells quantitatively regulate the onset of severe MP287 disease in mice, we adoptively transferred intermediate (3×10^6^) or low (1×10^6^) numbers of CD4^+^ T cells into MP287-infected CD4-deficient mice (*Cd4*^-/-^) (**Figures 1G and S1F-G**). The numbers of CD4^+^ T cells are intermediate or low in comparison to the infected WT mice. *Cd4*^-/-^ mice developed severe TB, while mice receiving intermediate numbers of CD4^+^ T cells lost more body weight than mice that received lower cell numbers; these mice were in fact protected against disease, controlling lung damage and bacterial load (**Figures 1H-J**). Lung weight and lung cell infiltrate were increased in *Cd4*^*-/-*^ mice and mice transferred with intermediate numbers of CD4^+^ T cells, in comparison to mice transferred with lower CD4^+^ T cell numbers (**Figures S1H-J**). Of note, TB in mice transferred with 1×10^6^ CD4^+^ T cells was much less severe than in WT mice. (**Figure 1A**). These results suggest that excessive accumulation of lung parenchymal CD4^+^ T cells contributes to the worsening of TB caused by hypervirulent mycobacteria.

### T-cell-specific P2RX7 expression promotes lung parenchymal CD4^+^ T cell accumulation in response to hypervirulent mycobacteria

After defining that excessive accumulation of lung parenchymal CD4^+^ T cells is associated with severe disease in response to MP287, we sought to define how CD4^+^ T cells form lung residency in this context. We have previously defined that the eATP receptor P2RX7 promotes the long-term residency of CD8^+^ T cells in non-lymphoid tissues (Borges da Silva et al., 2018; Borges da Silva et al., 2020), but whether the same is true for CD4^+^ T cells is less clear. We measured P2RX7 protein expression in lung intravascular and parenchymal CD4^+^ T cells in MP287-infected mice. P2RX7 was highly expressed in lung parenchymal CD4^+^ T cells of infected mice in comparison to intravascular counterparts (**Figure 2A**).

**Figure 2.**
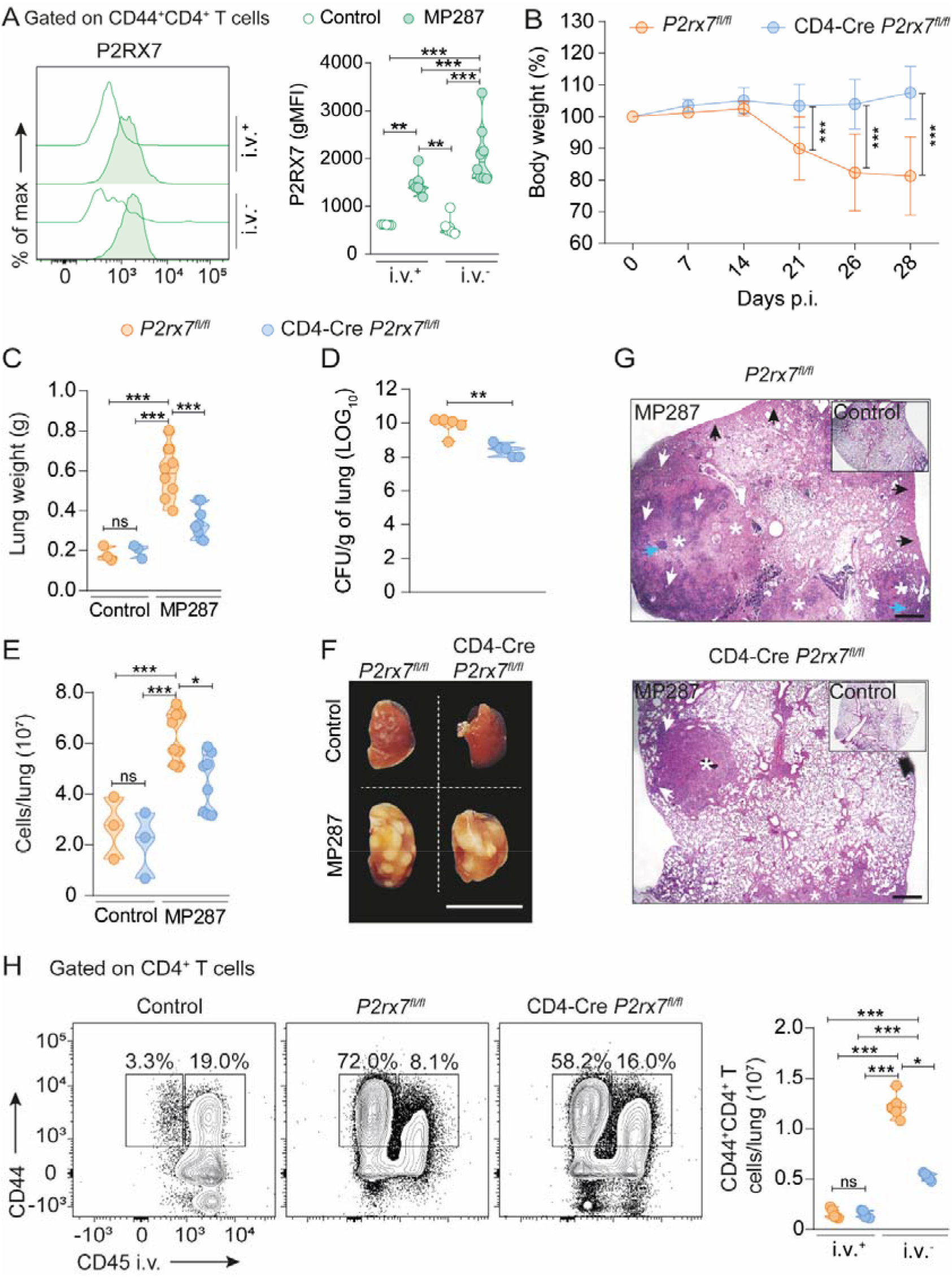
T-cell intrinsic P2RX7 promotes lung-parenchymal CD4^+^ T cell accumulation and severe TB caused by hypervirulent mycobacteria. (**A-H**) WT (C57BL/6 or *P2rx7*^*fl/fl*^) or T cell-P2RX7-KO (CD4-Cre *P2rx7*^*fl/fl*^) mice were infected with ∼100 CFU of the *Mbv*-MP287 strain. Non-infected mice were used as controls. At 28 days p.i., mice were injected i.v. with anti-CD45 fluorescent antibodies 3 minutes before lung harvest. (**A**) Histograms of P2RX7 expression in CD45iv^+^ and CD45iv^-^ CD44^+^CD4^+^ T cells of WT (C57BL/6) mice. The violin graphs show the geometric mean fluorescence intensity (gMFI) values of P2RX7 expression. (**B**) Percentages of body weights in relation to day 0. (**C**) Lung weight values. (**D**) CFU numbers per lung. (**E**) Total lung cell numbers. (**F**) Macroscopic images of right upper lung lobes (bar scales = 1 cm). (**G**) Representative lung sections stained with HE (bar scales = 500 μM). White asterisks indicate areas of necrosis, white arrows indicate area of alveolitis, blue arrows indicate bronchial obstruction and black arrows indicate perifocal exudative reaction. (**H**) Left: flow cytometry plots of CD44 and CD45i.v. expression in lung CD4^+^ T cells. Right: violin plots of the numbers of CD45iv^+^ and CD45iv^-^ CD44^+^CD4^+^ T cells per lung. Data are from 2 independent experiments; n = 3-5 per experimental group per experiment. Graphical data is shown as means ± SD. p values from One-way ANOVA (Tukey post-tests) (**A, C, E** and **H**) or unpaired T-tests (**B** and **D**) are indicated by ^*^, p <0.05; ^**^, p<0.01; ^***^, p <0.001.

Next, we investigated the role of P2RX7 in the lung parenchymal and intravascular CD4^+^ T cell responses to severe TB in WT and P2RX7-KO (*P2rx7*^*-/-*^) mice. Confirming previous studies (Amaral et al., 2014; Bomfim et al., 2017), P2RX7-KO mice were more resistant to MP287 infection than WT mice (**Figures S2A-C**). At 28 days p.i., numbers of lung parenchymal CD4^+^ T cells were reduced in infected *P2rx7*^*-/-*^ mice compared to infected WT mice, while comparable numbers were observed in the lung vasculature (**Figures S2D-E**).

Despite the increased expression of P2RX7 in lung CD4^+^ T cells of WT MP287-infected WT mice (**Figure 2A**), it is possible that the role of this receptor for lung CD4^+^ T cell accumulation in response to severe TB can be due to signaling in lung-infiltrating innate immune cells, which also express P2RX7 (Amaral et al., 2014; Bomfim et al., 2017). To evaluate whether the role of P2RX7 for lung-parenchymal CD4^+^ T cell accumulation is T cell-intrinsic or not, we used a T cell-specific P2RX7-KO (T cell-P2RX7-KO; CD4-cre*P2rx7*^*fl/fl*^) model. After MP287 infection, T cell-P2RX7-KO mice did not show the body weight loss observed in WT mice (*P2rx7*^*fl/fl*^) (**Figure 2B**). Moreover, T cell-P2RX7-KO mice had decreased lung weights in response to MP287 (**Figure 2C**) and better control of pulmonary mycobacterial growth (**Figure 2D**). Total lung cell numbers were also decreased in T cell-P2RX7-KO mice (**Figure 2E**), which was associated with diminished lung inflammation and necrosis (**Figures 2F-G**). Finally, T cell-P2RX7-KO mice had significantly decreased numbers of lung parenchymal CD4^+^ T cells in comparison to WT mice (**Figure 2H**). Together, these data indicate that T cell-intrinsic P2RX7 promotes the accumulation lung-parenchymal CD4^+^ T cells and subsequent severe TB in response to hypervirulent mycobacteria.

### P2RX7 is required for the lung parenchymal CD4^+^ T cells to protect against hypervirulent mycobacteria when in low numbers

We then investigated whether the previously observed protective role of restricted accumulation of lung-parenchymal CD4^+^ T cells (**Figures 1G-J**) is controlled by P2RX7. We adoptively transferred 1×10^6^ P2RX7-KO CD4^+^ T cells into MP287-infected CD4-KO hosts (*P2rx7*^*-/-*^*>Cd4*^*-/-*^) and compared them to CD4-KO mice transferred with WT CD4^+^ T cells (*WT>Cd4*^*-/-*^) (**Figure 3A**). *P2rx7*^-/-^>*Cd4*^-/-^ and non-transferred *Cd4*^*-/-*^ mice progressively lost body weight from day 21 p.i., which was not observed in infected WT>*Cd4*^-/-^ mice (**Figure 3B**). Likewise, we found increased lung weights in *P2rx7*^-/-^>*Cd4*^-/-^ mice compared to WT>*Cd4*^-/-^ mice (**Figure 3C**). Supporting a crucial role for the P2RX7 in CD4^+^ T cell effector function, higher bacterial loads were observed in the lungs of *P2rx7*^-/-^>*Cd4*^-/-^ mice compared to WT>*Cd4*^-/-^ mice (**Figure 3D**). We also found numerous granulomatous lesions and necrotic areas in infected *P2rx7*^-/-^>*Cd4*^-/-^ mice, while few areas of inflammation without necrosis were observed in WT>*Cd4*^-/-^ mice (**Figures 3E-F)**.

**Figure 3.**
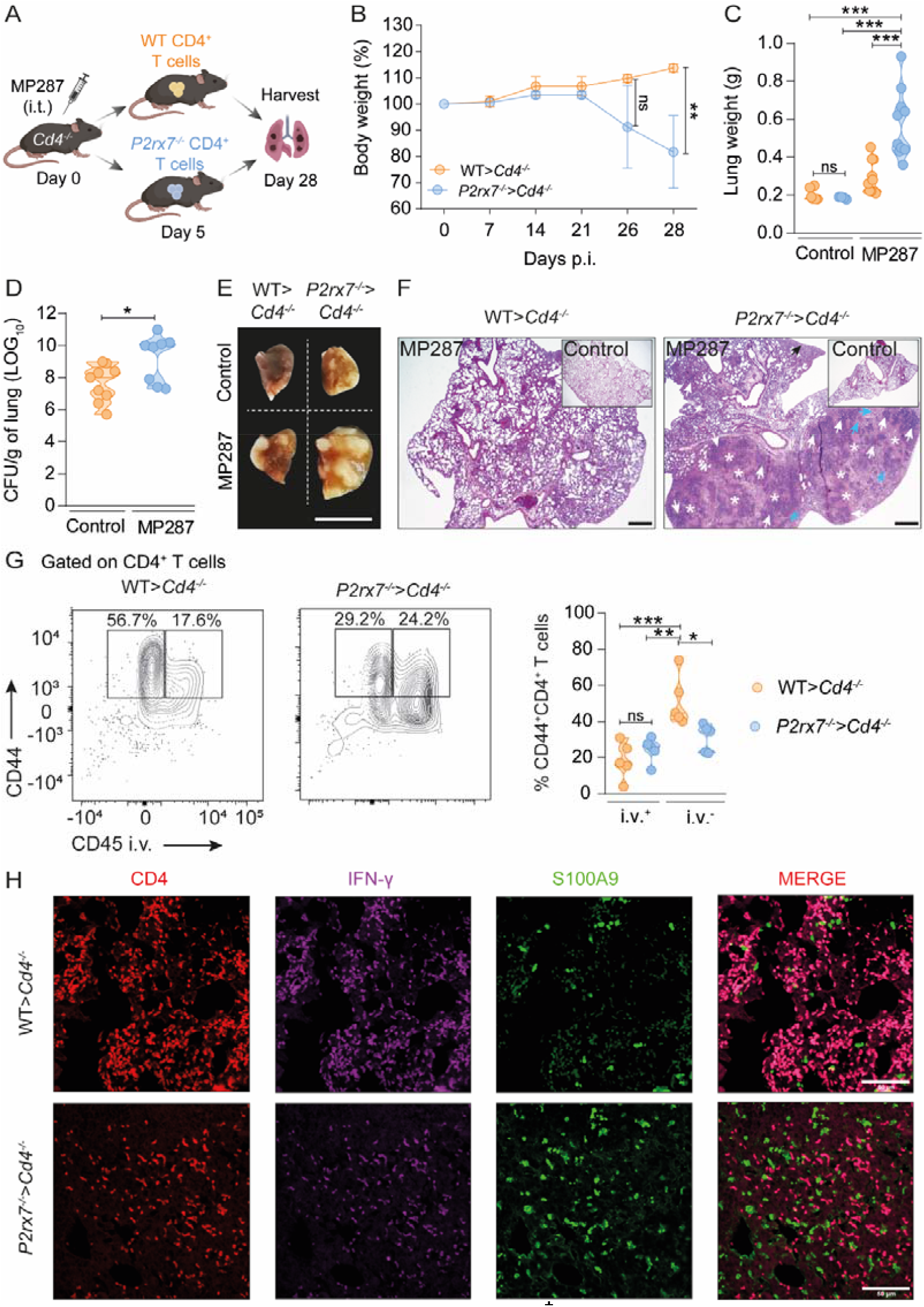
Restricted numbers of P2RX7-KO CD4^+^ T cells fail to transfer protection against hypervirulent mycobacteria. CD4-KO (*Cd4*^*-/-*^) mice were infected with ∼100 CFU of the *Mbv*-MP287 strain. Non-infected and infected *Cd4*^*-/-*^ mice were used as controls. At day 5 p.i. mice received splenic CD4^+^ T cells (1×10^6^) from WT and P2RX7-KO (*P2rx7*^*-/-*^) mice. At 28 days p.i., mice were injected i.v. with anti-CD45 fluorescent antibodies 3 minutes before lung harvest. (**A**) Schematic illustration of experimental protocol (Created with BioRender.com). (**B**) Percentages of body weights in relation to day 0. (**C**) Lung weight values. (**D**) CFU numbers per lung. (**E**) Macroscopic images of right upper lung lobes (bar scales = 1 cm). (**F**) Representative lung sections stained with HE (bar scales = 500 μm). White asterisks indicate areas of necrosis, white arrows indicate area of alveolitis, blue arrows indicate bronchial obstruction and black arrows indicate perifocal exudative reaction. (**G**) Left: flow cytometry plots of CD44 and CD45i.v. expression on CD4^+^ T cells. Right: frequencies (%) of CD45i.v.^+^ and CD45i.v.^-^ cells in CD44^+^CD4^+^ T cells. (**H**) Representative confocal images of CD4, IFN-γ and S100A9 expression in infected lung tissue (bar scale = 50 μm). The cells were stained for CD4 (red), IFN-γ (magenta) and S100A9 (green). Data are from 2–4 independent experiments; n = 3-5 per experimental group per experiment. Graphical data is shown as means ± SD p values from One-way ANOVA (Tukey post-tests) (**C, D** and **G**) or unpaired T-tests (**B**) are indicated by ^*^, p <0.05; ^**^, p<0.01; ^***^, p <0.001.

To determine how low numbers of P2RX7-KO CD4^+^ T cells failed to transfer protection to CD4-KO mice, CD4^+^ T cells in the lung parenchyma and vasculature of WT>*Cd4*^-/-^and *P2rx7*^-/-^>*Cd4*^-/-^ mice were analyzed. While most lung WT CD4^+^ T cells were found in the lung parenchyma of infected mice, P2RX7-KO CD4^+^ T cells were predominantly associated with the lung vasculature and excluded from the parenchyma (**Figure 3G**). Moreover, fewer IFN-γ-producing CD4^+^ T cells infiltrated the lungs of infected *P2rx7*^-/-^>*Cd4*^-/-^ mice compared to infected WT>*Cd4*^-/-^ mice (**Figure 3H**). In addition, the staining for S100A9 alarmin evidenced increased lung tissue damage in infected *P2rx7*^-/-^>*Cd4*^-/-^ mice. These results demonstrate that CD4^+^ T cell-intrinsic P2RX7, despite promoting lung damage when high CD4^+^ T cell numbers are present in the lung parenchyma, is paradoxically required for restricted numbers of CD4^+^ T cells to protect against MP287 infection.

### P2RX7 promotes lung CD4^+^ T cell accumulation by controlling *in-situ* proliferation in response to severe TB

To evaluate how WT and P2RX7-KO CD4^+^ T cells respond to MP287 infection in a competitive environment, WT (CD45.1^+^) and P2RX7-KO (CD45.2^+^) CD4^+^ T cells were co-transferred to *Cd4*^*-/-*^ mice at 5 days p.i. (**Figure 4A**). In these experiments, nanobody-mediated ARTC2.2 blockade was used to prevent the death of parenchymal T cells via P2RX7 (Borges da Silva et al., 2019). On day 28 p.i., WT CD4^+^ T cells accumulated mainly in the lung parenchyma, whereas P2RX7-KO CD4^+^ T cells were mostly restricted to the lung vasculature (**Figures 4B-C**). No difference was observed in the frequencies of WT and P2RX7-KO parenchymal IFN-γ - producing cells, but the absolute numbers of this population were higher among WT cells (**Figures 4D-E**). Additionally, there was no difference in the frequencies of T-bet^+^ CD4^+^ T cells (**Figure 4F**).

**Figure 4.**
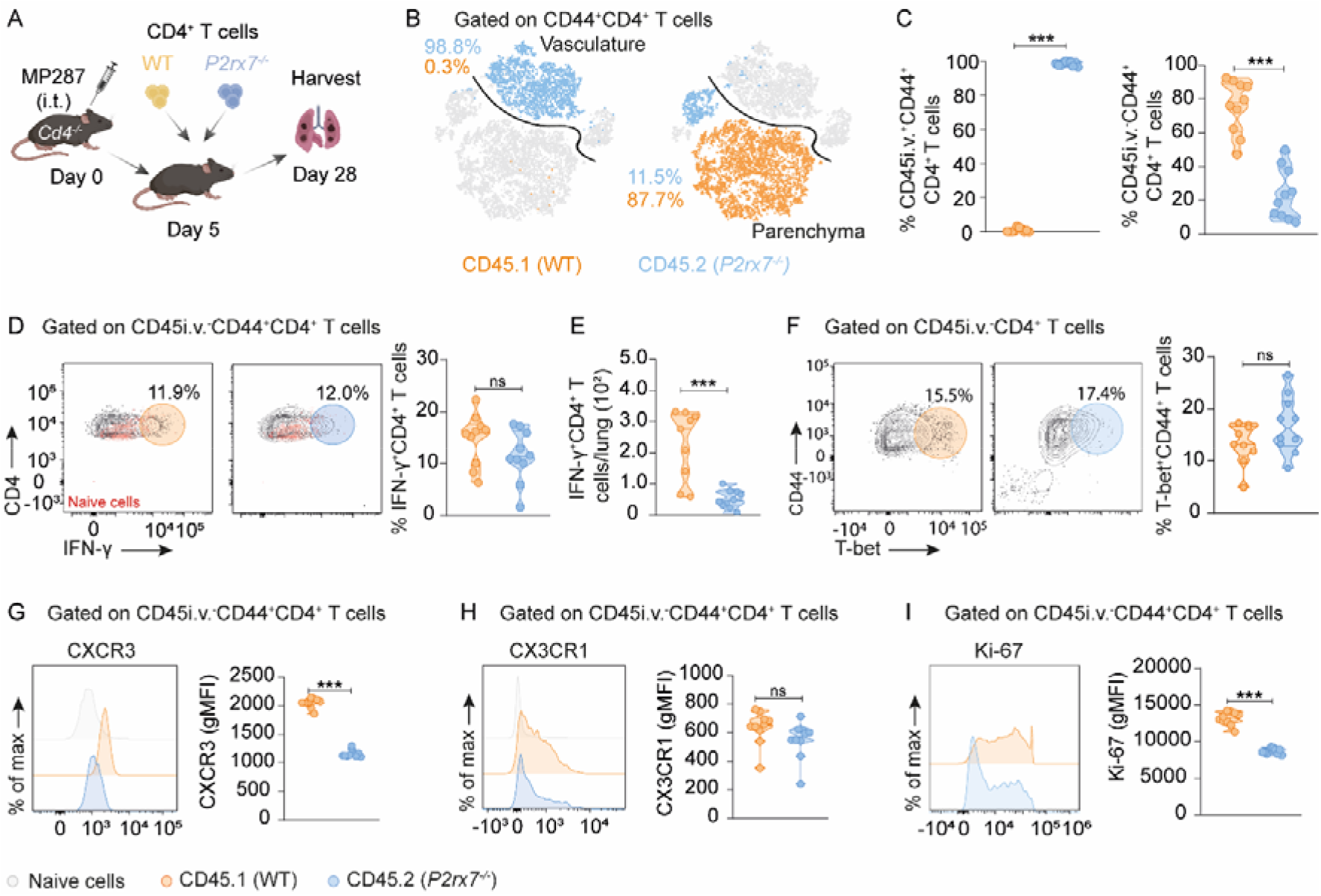
Cell-intrinsic P2RX7 expression is required for CD4^+^ T cell lung parenchymal establishment in response to severe TB. CD4-KO (*Cd4*^*-/-*^) mice were infected with ∼100 CFU of the *Mbv*-MP287 strain. On day 5 p.i. splenic CD4^+^ T cells (1×10^6^) from WT (CD45.1^+^) and P2RX7-KO (CD45.2^+^; *P2rx7*^*-/-*^) mice were co-transferred (1:1) to infected *Cd4*^*-/-*^ mice. At 28 days p.i., mice were injected i.v. with anti-CD45 fluorescent antibodies 3 minutes before lung harvest. (**A**) Schematic illustration of the experimental protocol (Created with BioRender.com). (**B**) T-distributed stochastic neighbor embedding (t-SNE) maps of transferred CD4^+^ T cells in the lung vasculature and parenchyma. (**C**) Frequencies (%) of CD45i.v.^+^ and CD45i.v.^-^ CD44^+^CD4^+^ T cells. (**D**) Left: flow cytometry plots of IFN-γ and CD4 expression in CD45i.v.^-^CD44^+^CD4^+^ T cells. Right: frequencies (%) of IFN-γ^+^ cells. (**E**) Numbers of IFN-γ ^+^CD45i.v.^-^CD44^+^CD4^+^ T cells per lung. (**F**) Left: flow cytometry plots of CD44 and T-bet expression in CD45i.v.^-^CD4^+^ T cells. Right: frequencies (%) of T-bet^+^ cells. (**G-I**) Histograms and gMFI values of CXCR3, CX3CR1 and Ki-67 expression in CD45i.v.^-^ CD44^+^CD4^+^ T cells. Naive CD4^+^ T cells were analyzed as a negative control. Data are from 2 independent experiments; n=5 per experimental group per experiment. Graphical data is shown as means ± SD. p values from unpaired T-test are indicated by ^*^, p <0.05; ^**^, p<0.01; ^***^, p <0.001.

We then evaluated the expression of molecules associated with the establishment of CD4^+^ T cells in the lung parenchyma (residency) or vasculature. In co-transferred and infected *Cd4*^*-/-*^ mice, WT CD4^+^ T cells expressed higher levels of CXCR3 and KLRG1 compared to P2RX7-KO CD4^+^ T cells (**Figure 4G and S3A**). The CX3CR1 was expressed at the same levels on WT and P2RX7-KO CD4^+^ T cells (**Figure 4H**). In contrast, CD69 expression was up-regulated in P2RX7-KO CD4^+^ T cells compared to CD4^+^ WT cells (**Figure S3B**). We did not compare the expression of these molecules in the intravascular compartment due to low numbers of WT CD4^+^ T cells. Overall, these results show that increased lung parenchymal residency is associated with CXCR3 expression in WT CD4^+^ T cells.

We also assessed the *in-situ* proliferation rates of parenchymal CD4^+^ T cells. This analysis showed that P2RX7-KO CD4^+^ T cells have impaired ability to proliferate in the lung parenchyma (**Figure 4I**). To confirm this result, CD4^+^ T cells were isolated from the lungs of infected mice for *ex vivo* proliferation assessment. Like our previous data, WT CD4^+^ T cells proliferated more efficiently than P2RX7-KO CD4^+^ T cells in response to MP287 (**Figure S3C**). These results show that, in response to hypervirulent TB, cell-intrinsic P2RX7 expression determines the ability of CD4^+^ T cells to express CXCR3, to establish residency in lung parenchyma and to proliferate *in-situ*.

### P2RX7 is not required for the activation and differentiation of effector CD4^+^ T cells in lung-draining lymph nodes of MP287-infected mice

To investigate whether the effects of P2RX7 on CD4^+^ T cells were determined prior to lung infiltration, we analyzed WT and P2RX7-KO CD4^+^ T cells from the mediastinal lymph nodes (medLNs) of co-transferred, MP287-infected *Cd4*^*-/-*^ mice (**Figure S4A**). The frequency of experienced (CD44^+^) P2RX7-KO CD4^+^ T cells is higher compared to WT CD4^+^ T cells in the medLNs (**Figure S4B**). Increased frequencies of WT T-bet^+^ CD4^+^ T cells were observed (**Figure S4C**). WT and P2RX7-KO CD4^+^ T cells co-expressed CXCR3 and CX3CR1 (**Figure S4D**).

However, in contrast to what was observed in the lung, P2RX7-KO CD4^+^ T cells expressed higher levels of CXCR3 and CX3CR1 than WT CD4^+^ T cells in medLNs (**Figures S4E and S4F**), making it unlikely that their lung phenotypes were acquired during early activation. These results demonstrate that both WT CD4^+^ T cells and P2RX7-KO CD4^+^ T cells leave medLNs with the ability to localize either in the vasculature or in the lung parenchyma. Therefore, they indicate that the eATP-P2RX7 interaction in the lung determines the fate of these cells during severe TB.

### T-cell intrinsic P2RX7 is crucial for lung inflammation and the establishment of lung parenchymal CD4^+^ T cells during Influenza infection

We then tested if the role attributed to CD4-specific P2RX7 in response to hypervirulent mycobacteria is infection-specific or reflects a phenomenon generalizable to other lung infections. We used viral infection by the PR8 strain of Influenza virus. Like what we observed in response to MP287, T cell-P2RX7-KO mice infected with the PR8 strain lost less body weight than WT mice (**Figure 5A**). T cell-P2RX7-KO mice also had decreased lung weights compared to WT mice on the day 14 p.i. with influenza (**Figure 5B**). On day 7 p.i., the viral load was higher in the lungs of WT mice than in those of P2RX7-KO mice, but at day 14 p.i. this difference was no longer observed (**Figure 5C**). Lower cell infiltration was also observed in T cell-P2RX7-KO mice (**Figure 5D**), which paralleled smaller lung sizes in comparison to WT mice (**Figures 5B and 5E**). This indicates a faster recovery of lung tissue. Indeed, T cell-P2RX7-KO mice displayed reduced lung inflammation and fibrosis after virus elimination (**Figure 5F**). We then assessed the location of CD4^+^ T cells in PR8-infected lungs. WT mice had a preferential accumulation of lung parenchymal CD44^+^CD4^+^ T cells, while T cell-P2RX7-KO showed a decrease in the number of parenchymal CD4^+^ T cells (**Figure 5G**).

**Figure 5.**
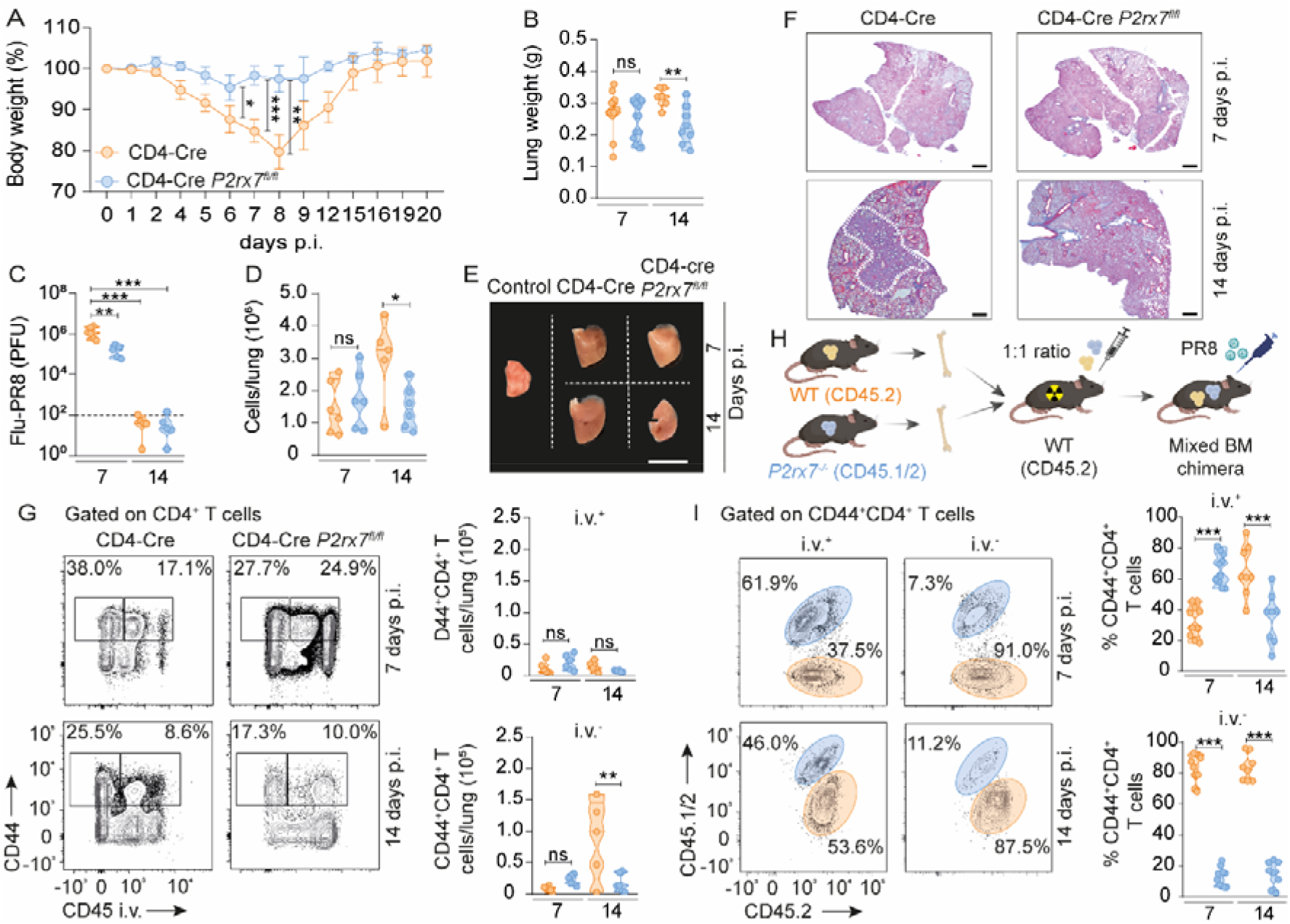
T-cell intrinsic P2RX7 promotes the accumulation of lung parenchymal CD4^+^ T cells and lung inflammation in response to influenza virus. (**A-G**) WT (CD4-Cre) and T cell-P2RX7-KO (CD4-Cre *P2rx7*^*fl/fl*^) mice were infected with Influenza virus PR8 strain (∼1400 PFU). At 7 and 14 days p.i., mice were injected i.v. with anti-CD45 fluorescent antibodies 3 minutes before lung harvest. (**A**) Percentages of body weights in relation to day 0. (**B**) Lung weight values. (**C**) Viral loads. (**D**) Total lung cell numbers. (**E**) Macroscopic images of right upper lung lobes (bar scales = 1 cm). (**F**) Representative lung sections stained with Masson’s trichrome (bar scales = 20 μM). (**G**) Left: low cytometry plots of CD44 and CD45 i.v. expression on CD4^+^ T cells. Right: The violin plots show the number of CD45i.v.^+^ and CD45i.v.^-^ CD44^+^CD4^+^ T cells. (**H-I**) In some experiments, WT (CD45.1^+^) mice were lethally irradiated and reconstituted with a 1:1 mix of WT (CD45.2^+^) and T cell-P2RX7-KO (CD45.1/2^+^) bone marrow cells. After 4-8 weeks, bone marrow chimeric mice were infected with PR8. (**H**) Schematic illustration of the experimental protocol (Created with BioRender.com). (**I**) Left: flow cytometry plots of CD45.1 and CD45.2 expression on CD45i.v.^+^ and CD45i.v.^-^ CD44^+^CD4^+^ T cells. Right: frequencies (%) of CD45.1^+^ and CD45.2^+^ cells. Data are from 2 independent experiments; n = 3-5 per experimental group per experiment. Graphical data is shown as means ± SD. p values from One-way ANOVA (Tukey post-tests) are indicated by ^*^, p <0.05; **, p<0.01; ^***^, p <0.001.

P2RX7 is crucial for the survival of regulatory T cells (Tregs), thus we analyzed the number of Foxp3^+^ Tregs in the lung parenchyma (**Figure S5A**). The number of Tregs was not different between WT and T cell-P2RX7-KO mice in hypervirulent mycobacterial infection. During PR8 infection, Treg numbers were higher in WT mice at day 7 p.i., with a similar trend (but not significant) at day 14 p.i. (**Figure S4A**). These results make it unlikely that the decreased lung pathology observed in T cell-P2RX7-KO mice is due to increased accumulation of lung Tregs.

To test the role of P2RX7 for lung CD4^+^ T cell accumulation in response to influenza in a competitive setup, mixed bone marrow chimera mice were generated and infected with PR8 (**Figures 5H and S5B**) (Borges da Silva et al., 2018). On days 7 and 14 p.i., the lung parenchyma of infected chimeric mice was enriched with WT CD4^+^ T cells in comparison to P2RX7-KO CD4^+^ T cells (**Figure 5I**). Together, these data demonstrate that P2RX7 is crucial for the exacerbation of lung inflammation mediated by parenchymal CD4^+^ T cell infiltration in response to influenza.

### P2RX7 in lung parenchymal CD4^+^ T cells increases the expression of genes and proteins associated with cell migration and proliferation in response to influenza

Next, we sought to define the molecular pathways differentially expressed between lung-parenchymal WT and T cell-P2RX7-KO CD4^+^ T cells. We sorted lung CD45i.v.^-^, lung CD45i.v.^+^ and spleen CD4^+^ T cells in mice infected with PR8 and performed RNA-seq analysis. From this, we detected >1000 differentially expressed genes between WT and T cell-P2RX7-KO cells in lung i.v.^+^ and lung i.v.^-^ compartments, and less than 200 between spleen CD4^+^ T cell populations (**Figures 6A and S6A**). The few forming lung-parenchymal P2RX7-KO CD4^+^ T cells clustered apart from lung-parenchymal WT CD4^+^ T cells (**Figure 6B**). Unexpectedly, lung vascular-associated P2RX7-KO CD4^+^ T cells were transcriptionally similar to lung WT CD4^+^ T cells, vascular or parenchyma-associated. Both trends were also apparent when the top differentially expressed genes were assessed between these groups (**Figures 6C and S6B**).

**Figure 6.**
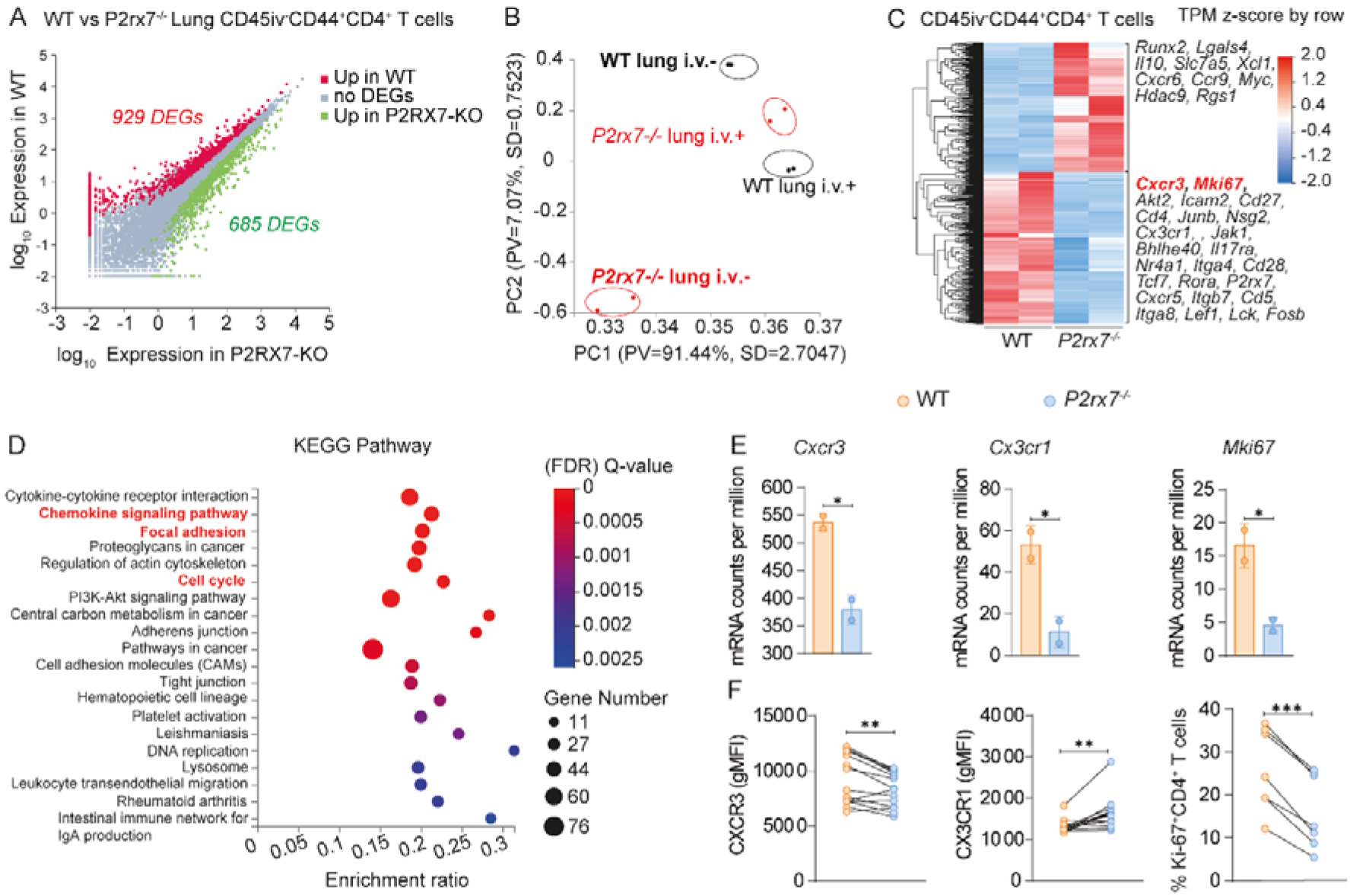
T cell-intrinsic P2RX7 induces expression of cell proliferation and tissue adhesion molecules in lung-parenchymal CD4^+^ T cells. (**A-E**) WT (CD4-Cre) and T cell-P2RX7-KO (CD4-Cre *P2rx7*^fl/fl^) mice were infected with influenza (PR8, ~1400 PFU). At day 7 p.i., lung CD45i.v.^-^, lung CD45i.v.^+^ and spleen CD44^+^ CD4^+^ T cells were sorted and RNA-seq analysis was performed. (**A**) Scatter plot with differentially expressed genes (DEGs) highlighted between WT and P2RX7-KO lung CD45i.v.^-^CD4^+^ T cells. (**B**) Principal Component Analysis (PCA) plots representing the relative transcriptional signatures of lung CD45i.v.^-^ and CD45i.v.^+^ CD4^+^ T cells. (**C**) Heatmap displaying the DEGs between WT and P2RX7-KO lung CD45i.v.^-^CD4^+^ T cells; representative DEGs higher on WT or P2RX7-KO CD4^+^ T cells are shown. (**D**) List of transcriptional pathways enriched in lung WT CD45i.v.^-^ CD4^+^ T cells in comparison to P2RX7-KO counterparts. (**E**) Values for mRNA counts per million for *Cxcr3, Cx3cr1, Cd69* and *Mki67* in lung WT and P2RX7-KO CD45i.v.^-^CD4^+^ T cells. (**F**) In other experiments, WT and T cell-P2RX7-KO mice were infected with PR8 and protein expression of different molecules was assessed by flow cytometry (day 7 post-infection). The gMFI values for CXCR3, CX3CR1, CD69 and percentages of Ki-67^+^ cells in lung WT and P2RX7-KO CD45i.v.^-^CD4^+^ T cells.

We also performed pathway enrichment analyses of these experimental groups. In comparison to lung parenchymal P2RX7-KO, WT CD4^+^ T counterparts had increased expression of (among other pathways) cell cycle, adhesion and chemokine signaling pathway genes (**Figure 6D**). Confirming these trends, the expression of the proliferation gene *Mki67* (which encodes Ki-67), as well as of *Cxcr3* and *Cx3cr1*, was decreased in lung-parenchymal P2RX7-KO cells (**Figure 6E**). Protein expression of Ki-67 and CXCR3 was correspondently diminished in lung-parenchymal T cell-P2RX7-KO CD4^+^ T cells, while expression of CX3CR1 was increased (**Figure 6F**). These results evidence that, in response to influenza, P2RX7 expression leads to modifications in lung parenchymal CD4^+^ T cells at the transcriptional and protein levels. Among these changes, increases in expression of the lung-homing chemokine receptor CXCR3 and of in situ proliferation (both also observed in response to *Mbv*-MP287 infection) suggest that some of these P2RX7-induced CD4^+^ T cell adaptations in the lung tissue are common to multiple lung infections.

### The accumulation of lung parenchymal CD4^+^ T cells depends on CXCR3 in influenza

We then evaluated whether CXCR3 is critical for the accumulation of lung parenchymal CD4^+^ T cells in influenza, as the need for this chemokine receptor may explain the requirement of P2RX7 in this process. PR8-infected WT and P2RX7-KO mice were treated with anti-CXCR3 blocking antibodies during the acute phase of infection (**Figure 7A**). This treatment reduced the lung parenchymal CD4^+^ T cell population; anti-CXCR3-treated WT mice had similar numbers of lung parenchymal CD4^+^ T cells than treated or untreated T cell-P2RX7-KO mice (**Figure 7B**). These results indicate that CD4^+^ T cells depend on the expression of CXCR3 to accumulate in the lung parenchyma and P2RX7 upregulates this chemokine receptor.

**Figure 7.**
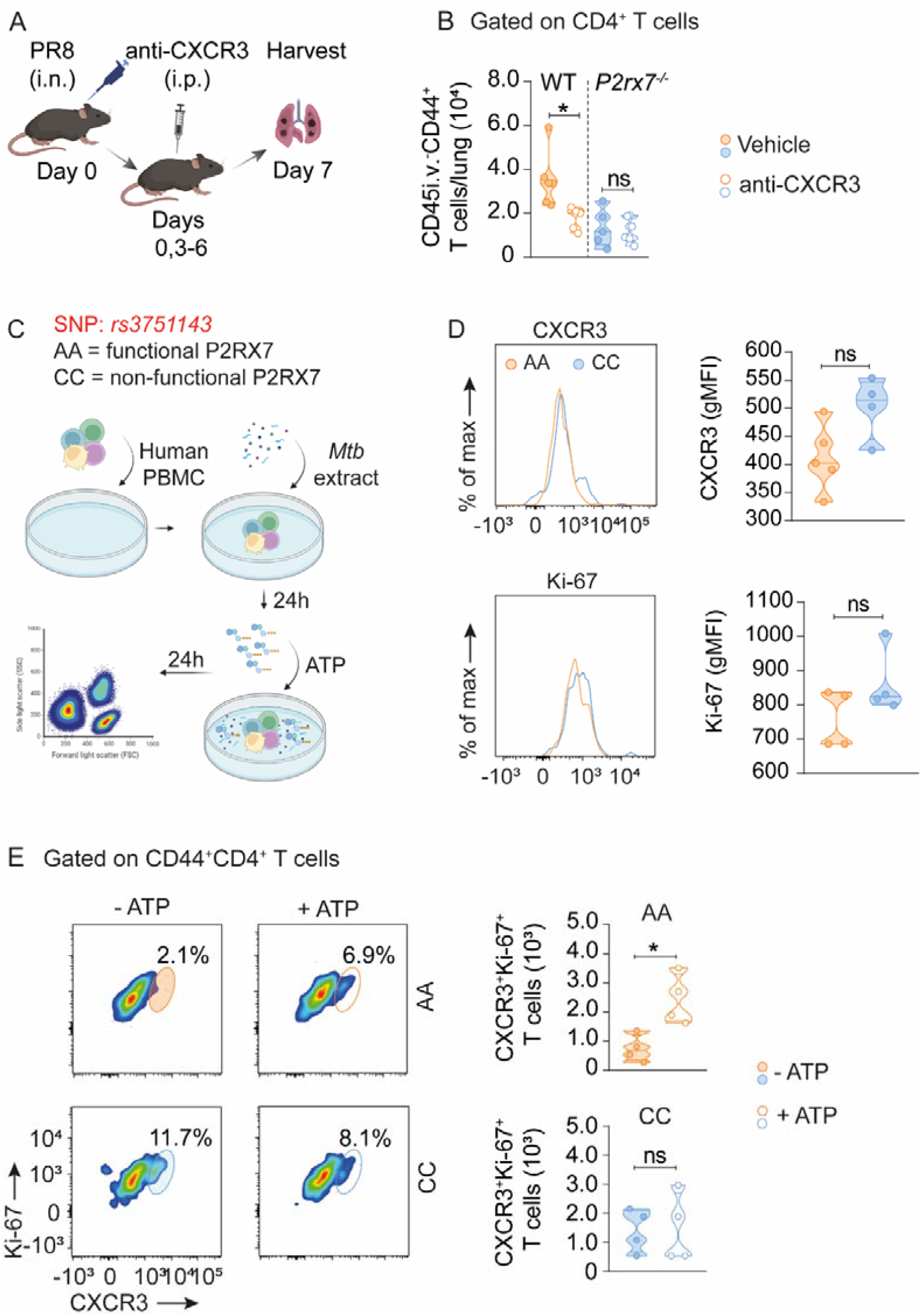
CXCR3 is critical for lung CD4^+^ T cell accumulation in influenza and is upregulated by P2RX7 activation in human CD4^+^ T cells that respond to mycobateria. (**A-B**) WT and *P2rx7*^*-/-*^ mice were infected with influenza (PR8, ~1400 PFU) and treated intraperitonially with anti-CXCR3 antibody on days 0, 3, 4 and 6 of infection. At 7 days p.i., mice were injected i.v. with anti-CD45 fluorescent antibodies 3 minutes before lung harvest. (**A**) Schematic illustration of the experimental protocol (Created with BioRender.com). (**B**) Violin plots of the numbers of CD45i.v.^-^CD44^+^CD4^+^ T cells per lung. (**C-E**) PBMCs (1×10^5^) from two patients with active leprosy, one of each carrying a loss-of-function P2RX7 SNP (*rs3751143 A>C*), were stimulated with a *Mtb* extract for 24h. Cells were then cultured with eATP (50 μM) for a further 24 h and analyzed by flow cytometry. (**C**) Schematic illustration of the experimental protocol (Created with BioRender.com). (**D**) Histograms and gMFI values of CXCR3 and Ki-67 expression in CD44^+^CD4^+^ T cells. (**E**) Right: flow cytometry plots of Ki-67 and CXCR3 expression in *Mtb* extract-stimulated CD4^+^ T cells cultured with or without eATP. Left: violin plots of the number of Ki-67^+^CXCR3^+^CD4^+^ T cells. Data are from 2 independent experiments; n = 2-4 per experimental group per experiment. Graphical data is shown as means ± SD p values from One-way ANOVA (Tukey post-tests) (**B**) or unpaired T-tests (**D and E**) are indicated by ^*^, p <0.05; ^**^, p<0.01; ^***^, p <0.001.

### P2RX7 activation directly induces CXCR3 upregulation in human CD4^+^ T cells that respond to mycobacteria

We also investigated whether P2RX7 activation upregulates CXCR3 expression and the proliferative response in PBMC CD4^+^ T cells from patients with the active form of leprosy, an infection caused by *Mycobacterium leprae*. This analysis was performed in a donor with a loss-of-function P2RX7 single-nucleotide polymorphism (SNP *rs3751143* CC - non-functional P2RX7) and a non-polymorphic donor (AA - functional P2RX7) (Souza et al., 2021). PBMCs were stimulated with a *Mtb* extract for 24 h and then cultured with ATP for a further 24 h in order to mimic the eATP-rich environment that cells find in the lung parenchyma (**Figure 7C**). No difference was observed in CXCR3 and Ki-6 expression before adding eATP to the culture (**Figure 7D**). Adding eATP to cell cultures increased the Ki-67^+^CD44^+^CD4^+^ T cell population for the AA donor, but not for the CC donor (**Figure 7E**). Corroborating the data we obtained in a murine model, these results indicate that P2RX7 activation by eATP upregulates CXCR3 expression and the proliferative response in human CD4^+^ T cells that respond to mycobacteria.

## DISCUSSION

CD4^+^ T cells are crucial for the defense of the host against pulmonary infections, being a featured vaccine target (Sant et al., 2018; Turner and Farber, 2014). However, there is a great dilemma involving the CD4^+^ T cell response during pulmonary infections (Barber, 2017). Despite being potent in providing protection, excessive effector CD4^+^ T cell responses can bring irreversible damage to tissues with low regenerative capacity (Moguche et al., 2017). Our present work suggests that the magnitude of lung CD4^+^ T cell accumulation determines the positive versus negative outcome of their effect in response to lung infections such as TB and influenza.

The hypervirulent MP287 mycobacteria has been shown to better mimic severe TB disease in humans (Amaral et al., 2014; Bomfim et al., 2017; Santiago-Carvalho et al., 2021). Differences between *Mbv*-MP287 and the standard *Mtb*-H37Rv are not limited to pathology induction: lung CD4^+^ T cell populations infiltrate and localize in different lung tissue compartments when comparing these two infections. In mice infected with H37Rv, *Mtb*-induced CD4^+^ T cells are located mostly in the pulmonary vasculature (Sakai et al., 2014) with low-to-intermediate numbers accumulating in the parenchyma. MP287 infection, in contrast, induces high numbers of lung parenchymal CD4^+^ T cells. These comparisons raised the question that the excessive accumulation of CD4^+^ T cells in the lung parenchyma could be the cause and not the consequence of the worsening of MP287 infection. CD4^+^ T cell depletion and adoptive transfer experiments indicated this is indeed the case. Moreover, these results evidenced that, even in the context of MP287 infection, low-to-intermediate accumulation of lung parenchymal CD4^+^ T cells is sufficient to promote protection. Therefore, our data support a working model where a goldilocks intensity of lung CD4^+^ T cell infiltration is necessary to protect against lung-tropic infections while avoiding excessive tissue damage. Mechanistically, this may be explained by differential production of CD4^+^ T cell-derived pro-inflammatory cytokines, such as IFN-γ or TNF-α. High levels of these cytokines can promote exacerbated recruitment and *in situ* activation of innate immune cells, exaggerated host cell death or increased pulmonary fibrosis (Amaral et al., 2014; Hillaire et al., 2013; Hirahara et al., 2021). In contrast, the entry of low-to-intermediate numbers of CD4^+^ T cells would lead to levels of IFN-γ and TNF-α sufficient to promote the elimination of viruses or bacteria, while off-target effects in the lung tissue itself being limited.

P2RX7 was defined by us to play a crucial role in the generation of CD8^+^ T^RM^ cells (Borges da Silva et al., 2018; Borges da Silva et al., 2020), and CD4^+^ T_RM_ cells in several non-lymphoid tissues reportedly express high levels of P2RX7 (Beura et al., 2019), including a subset of lung CD4^+^ T_RM_ cells observed long after elimination of influenza (Son et al., 2021; Swarnalekha et al., 2021). To our knowledge, however, our work is the first to demonstrate that expression of P2RX7 is required for the establishment of infection-induced CD4^+^ T cells in the lung parenchyma. P2RX7 is a low affinity eATP receptor (Di Virgilio et al., 2017), therefore its action has been associated with eATP accumulation induced by tissue cell death during infections (Amaral et al., 2014; Borges da Silva et al., 2020; Di Virgilio et al., 2017). Multiple immune cells and lung epithelial cells express P2RX7, and our group has previously shown that pharmacological P2RX7 inhibition led to decreased CD4^+^ T cell numbers in the lung parenchyma in response to MP287 (Santiago-Carvalho et al., 2021). Our results expand these findings by identifying that the infection-induced accumulation of lung parenchymal CD4^+^ T cells depends on P2RX7 signaling directly on T cells. The P2RX7-mediated increase in lung parenchymal CD4^+^ T cells is also a causal factor of lung damage and development of severe TB-MP287 or influenza-PR8 infections. This was evidenced by the phenotype of infected T cell-P2RX7-KO mice, which are protected from weight loss and lung inflammation. On the flip side, transfer of limited numbers of P2RX7-KO CD4^+^ T cells was insufficient to transfer protection against MP287 infection. This further suggests the diminished ability of P2RX7-KO CD4^+^ T cells to enter the lung parenchyma. Moreover, it indicates that P2RX7, by decreasing the CD4-intrinsic threshold for lung tissue infiltration, is important to define whether CD4^+^ T cell responses to lung infections will be beneficial or deleterious.

It is likely that not only P2RX7 is a controlling factor for that, but also the abundance of eATP. In the peak of MP287 infection, eATP levels are likely higher in the lung parenchyma (Amaral et al., 2014; Bomfim et al., 2017). In future studies, it will be important to evaluate whether the P2RX7-eATP induction of lung CD4^+^ T cell infiltration is a continuum, and what are the respective roles of overall lung pathogen load versus infection of specific lung cell types. On the latter, it is good to remember that influenza and mycobacteria infect different lung cell types (O’Garra et al., 2013; Thomas et al., 2006) and induce distinct inflammation modalities (Damjanovic et al., 2011; Nunes-Alves et al., 2014), yet the role of CD4^+^ T cell-specific P2RX7 is similar. This would suggest, instead, that lung pathogen load is the decisive factor for exacerbated eATP realease in this tissue.

We have previously identified that P2RX7 promotes the tissue residency of CD8^+^ T cells through the TGF-β signaling pathway (Borges da Silva et al., 2020). However, CD4^+^ T cells residing in non-lymphoid tissues do not express TGF-β-dependent signaling network molecules and form independently of TGF-βRII (Fonseca et al., 2022). Likewise, CD4^+^ T_RM_ cells do not express the TGF-β-induced, E-cadherin-binding molecule CD103 (Beura et al., 2019). Thus, the mechβanisms behind P2RX7-induced CD4^+^ T cell lung parenchymal establishment were unclear to us. In contrast to TGF-βRII and CD103, tissue-resident CD4^+^ T cells express the chemokine receptor CXCR3 (Beura et al., 2019). This is also true in response to mild *Mtb* infection, where CXCR3 is one of the main chemokine sensors determining the lung parenchymal homing of CD4^+^ T cells (Hoft et al., 2019). Our results in TB and influenza showed that the fewer lung parenchymal P2RX7-KO CD4^+^ T cells express lower levels of CXCR3 than WT conterparts, both at the protein and at transcriptional levels. Indeed, antibody-mediated blockade of CXCR3 was sufficient to reduce the accumulation of WT CD4^+^ T cells in lung parenchyma, with no effects on P2RX7-KO CD4^+^ T cells. Activation of this pathway possibly occurs only upon entry into the lung tissue. This is supported by our data showing that effector CD4^+^ T cells lacking P2RX7 exit draining lymph nodes expressing normal levels of CXCR3 and CX3CR1, which induce respectively parenchymal and intravascular T cell positioning (Gerlach et al., 2016; Sakai et al., 2014). Instead of a pre-determination of a tissue-parenchymal versus circulating fate during priming, our results suggest that sensing of lung eATP determines this fate, by inducing a further increase in CXCR3 expression. Of note, this was also observed by us in *Mtb*-responding human CD4^+^ T cells cultured *in vitro* with free ATP. It is still to be determined if lung tissue-derived eATP (and its interaction with P2RX7) directly alters the transcriptional and protein landscape of effector CD4^+^ T cells (including the enhance in CXCR3 expression), or if it merely serves as a *de facto* chemotactic signal to attract CD4^+^ T cells into the parenchyma, and indirectly regulating the tissue homing of TB- or influenza-induced CD4^+^ T cells. An indirect effect of eATP may also explain our results showing P2RX7-dependent *in situ* proliferation in lung parenchymal CD4^+^ T cells. Despite a previous report showing that eATP can induce the proliferation of human CD4^+^ T cells (Schenk et al., 2008), these findings can be confounded by the fact that another eATP receptor, P2RX4, can promote T cell activation during priming (Woehrle et al., 2010). Alternatively, the role of eATP-P2RX7 interaction as a chemoattractant for lung parenchyma may favor subsequent interactions of these lung CD4^+^ T cells with other tissue-localized mitogenic factors.

Regardless of these nuances, our work suggests a mechanism by which, in response to lung infections, P2RX7 favors lung CD4^+^ T cell parenchymal establishment via concomitant induction of CXCR3 high expression and of *in situ* proliferation. Again, the magnitude of this effect would be dependent on the P2RX7 expression level by effector CD4^+^ T cells, and on the amount of eATP released in the lung tissue. This may be, evidently, generalizable for tissue-resident CD4^+^ T cells induced by other infections or in other non-lymphoid tissues. Future research will be needed to test this hypothesis. Our group has already demonstrated that pharmacological blockade of P2RX7 partially reverts the exaggerated accumulation of parenchymal CD4^+^ T cells, preventing the generation of severe forms of TB (Santiago-Carvalho et al., 2021). Therefore, our study opens new perspectives for therapeutic interventions aimed at reducing the worsening of infectious diseases that attack the lung, without affecting the ability of the adaptive immune system to fight off these infections.

## Limitations of study

Infections caused by TB-MP287 and influenza-PR8 cannot be fully compared. Besides the distinct etiology of the respective infectious agents, MP287 infection generates 100% mortality within 30 days p.i. (Amaral et al., 2014; Bomfim et al., 2017), whereas PR8 viruses at the dose used here induces recovery of more than 50% of mice after acute infection (data not shown). These limitations may hinder our parallels traced here. Nevertheless, in both scenarios P2RX7 played a similar role in lowering the threshold for CD4^+^ T cell lung parenchymal accumulation. Moreover, previous reports had suggested a positive role for P2RX7 in the protection against milder forms of TB (Santos et al., 2013; Zheng et al., 2017), which may help solidify the notion that, more than species-specific nuances, it is the overall lung damage induced by pathogen load that dictates the role of CD4^+^ T cells and of P2RX7. More refined experimental tools to test these nuances will be subject of future research by our group. For this report, however, our goal was to determine the overall role of CD4^+^ T cell-specific P2RX7 in response to lung infections.

Another limitation we must consider is that our T cell-P2RX7-KO mouse model is also P2RX7-deficient on CD8^+^ T cells. We have reported that P2RX7 promotes CD8^+^ T_RM_ cells in response to systemic viruses (Borges da Silva et al., 2018; Borges da Silva et al., 2020), however in unpublished results we have not found, between WT and T cell-P2RX7-KO mice, major differences in the initial seeding of the lung parenchyma in response to influenza (data not shown), where these cells play a central role (Laidlaw et al., 2014). To mitigate this limitation, in several experiments we used adoptive transfer of CD4^+^ T cells into CD4-KO infected mice. This is an experimental tool our group has used before (Salles et al., 2017), to study the effects of P2RX7 deficiency only on CD4^+^ T cells. In this context, we chose low amounts of WT and P2RX7-KO CD4^+^ T cells for transfers, since we already know the pathological effect of high amounts of these cells in WT and P2RX7-KO mice. Nevertheless, in future studies, we aim to employ more refined experimental models to specifically address the role of CD4^+^ T cell-specific P2RX7 in the context of these lung infections.

## ACKNOWLEDGMENTS

We thank José Israel Lima, Silvana Silva, Maria Áurea, Juliana Azevedo and Verônica Lanes for technical assistance. This work was funded by Research Support of São Paulo (FAPESP) (M.R.D.L.: 2019/24700-8 and 2015/20432-8), National Council for Scientific and Technological Development (CNPq) (M.R.D.L.: 408909/2018-8 and 303810/2018-1) and the National Institute of Allergy and Infectious Diseases (NIAID) (H.BdS.: R00 AI139381, R01 AI170649).

## AUTHOR CONTRIBUTIONS

I.S.C., H.BdS. and M.R.D.L. conceived the project. I.S.C., H.BdS. and M.R.D.L. conceived the experiments. H.BdS, R.C.S., J.C.A.F., J.M.A.M., M.H.H. and E.L. provided mice or reagents for Biosafety Level 3 Laboratory experiments. I.S.C., G.A.S., C.C.B.B., F.M.A., B.M.G., T.V.K., M.V.P.C., S.V.D., B.M.M., R.S.d.N., E.L. and H.BdS. performed the experiments and analyzed the data. I.S.C., H.BdS. and M.R.D.L. wrote the manuscript.

## STAR METHODS

### Mice

Specific-pathogen-free (SPF) mice of of 6-8-week-old C57BL/6 (CD45.1 and CD45.2), *Cd4*^*-/-*^, *P2rx7*^*-/-*^, *P2rx7*^*flox/flox*^, CD4-Cre and CD4-Cre *P2rx7*^*flox/flox*^ strains were used. Male mice were used for experiments of severe tuberculosis and female mice for experiments with Influenza virus. The mice were kept in microisolators in the isogenic mouse facilities of the Institute of Biomedical Sciences of the University of São Paulo (USP) (São Paulo, Brazil), or of the Center for Biosciences and Biotechnology of the State University of Norte Fluminense (UENF) (Rio de Janeiro, Brazil) or from department of Immunology of Mayo Clinic (Arizona, USA) until the time of experimentation. To start the experiments, the mice destined to experiment with a severe tuberculosis model were transported to the Biosafety Level 3 (BSL-3) laboratory of the Institute of Pharmaceutical Sciences at USP or the Laboratory of Biology of Recognition at UENF, where all the experiments with the strain M. bovis MP287 were performed. Mice that were infected with the PR8 strain of Influenza virus were maintained in the Biosafety Level 2 (BSL-2) laboratory of the Mayo Clinic (Arizona, USA). In all experiments, mice were randomly assigned to experimental groups. All experimental procedures were approved by the institutional animal care and use committee of the institutions involved in the work (CEUA 5611150818, CEUA 402/2021 and IACUC A00005542-20).

#### Mycobacterium bovis (Mbv)

The clinical isolate *Mbv*-MP287/03 was provided by Dr. José Soares Ferreira Neto (School of Veterinary Medicine and Animal Science - University of São Paulo). The strain was kept and frozen at -80°C in Middlebrook 7H9 medium (Becton Dickinson, US) supplemented with ADC (albumin, dextrose and catalase) (Difco, US). Monitoring of mycobacterial growth was carried out by measuring Optical Density (OD 600nm) by spectrophotometry (Biochrom, model Libra S6). To carry out the experiments, the mycobacterial strain was removed from -80 ºC, thawed and added to Middlebrook 7H9 medium plus 10% of ADC enrichment medium. The suspension culture was vortexed (Biomatic, Brazil) and sonicated in an ultrasound bath (Ultrasonic Maxi cleaner 800 – Unique, Brazil) for 1 minute to disperse the lumps. The culture was incubated at 37 ºC and 5% of CO^2^ for 5-7 days.

#### Influenza virus

The Influenza virus PR8/34 strain was kindly provided by Dr. Jie Sun (University of Virginia). Aliquots containing ∼1.1×10^9^ PFU were stored in a freezer -80 ºC until the moment of infection.

### Infections

Infections with *Mbv*-MP287 were performed intratracheally (i.t.), by surgical incision, with ∼100 CFU in 90 μl of final volume. To be infected, the mice were anesthetized with a solution of ketamine (110mg/kg) and xylazine (15mg/kg), being inoculated from 80 to 100 μL (depending on the weight and sex of the animals) of anesthetic intraperitoneally. For Influenza virus infections the mice were anesthetized with isoflurane and 20 μl of viral inoculum (1400 PFU) was injected intranasally (i.n.).

### CD4^+^ T cells purification and adoptive transfer

Splenic CD4^+^ T cells were isolated from C57BL/6 and *P2rx7*^*-/-*^ mice by negative selection using the EasySep™ Mouse CD4^+^ T Cell Isolation Kit (STEMCELL), following the manufacturer’s protocol. After isolation, we obtained ∼94% purity. Next, CD4^+^ T cells (1×10^6^ or 3×10^6^) were transferred (1×10^6^ or 3×10^6^ per mouse) by i.v. to *Cd4*^*-/-*^ mice 5 days after infection with the MP287 Mbv strain.

### Intravascular labeling

For intravascular staining, mice were anesthetized and given intravenous (i.v.) injections of 2.5 μg of a fluorophore-labeled monoclonal antibody (mAb) against CD45 (30-F11) (BioLegend), and the lungs were harvested after 3 minutes, as described previously (Anderson et al., 2014).

### CD4^+^ T cell depletion

C57BL/6 mice were infected with the Mbv MP287 strain and on day 21 p.i. received a single i.p. of purified anti-α-CD4 antibody GK 1.5 (250 μg) (Bio X Cell). This day was chosen for the treatment, because it is when the aggravation of the disease in the mice begins. On day 28 p.i., mice were euthanized, and depletion was verified in the lung by flow cytometry (**Figure S1B**).

### Lung processing and cell preparation

The lung was collected, and the lobes were processed and digested with collagenase type IV (0.5 mg/mL) (Sigma-Aldrich) at 37º C for 40 minutes under agitation (200 rpm) (Amaral et al., 2019). The cell suspension obtained was homogenized and filtered through cell filters (Corning) and incubated with ACK Lysing Buffer (Thermo Fisher Scientific) at room temperature for one minute to deplete the erythrocytes. The lung cell suspensions were washed with PBS 10% fetal calf serum (Gibco, US). The cell suspension was centrifuged at 1,200 rpm for 5 minutes and resuspended in RPMI 1640 medium enriched with 10% FCS and 0.1% gentamicin (Gibco, US). Next, the viable lung cell numbers were determined using trypan blue exclusion assay and a hemocytometer.

### Flow cytometry analysis

T cells were isolated from the lung by tissue digestion with collagenase IV (0.5 mg/mL) and from the mediastinal lymph node (mLN) by mechanical digestion. In the adoptive cotransfer experiments using severe tuberculosis model and in the experiments with Influenza virus, 50 μg of Treg-Protector (anti-ARTC2.2) nanobodies (BioLegend) were injected i.v. 30 minutes before lung harvesting (Borges da Silva et al., 2019). Lung and mLN cells were stained using fluorochrome-labeled monoclonal antibodies to CD4, CD8, CD45, CD45.1, CD45.2, CD44, CD69, KLRG1, CX3CR1, CXCR3, IFN-γ, P2RX7, Ki-67 and T-bet (BD Biosciences). For ex vivo intracellular IFN-γ staining, lung cells were incubated with monensin (2 μM) for 5 hours at 37°C in 5% CO_2_ atmosphere, fixed, and permeabilized with BD cytofix/cytoperm kit (BD Biosciences) (Amaral et al., 2019). Live/dead dye was used to stain dead cells. I-A(b) Influenza A NP 311-325 (QVYSLIRPNENPAHK) major histocompatibility complex tetramers were produced in the National Institute of Allergy and Infectious Diseases Tetramer Core Facility (Emory University, Atlanta, GA). Cells were analysed using a LSRFortessa™, FACSymphony™ A5 flow cytometer (BD Biosciences). Conventional and spatial distribution by t-distributed stochastic neighbor embedding analysis (tSNE) was performed using FlowJo software (BD Biosciences) (Barbosa Bomfim et al., 2021).

### Histological analysis

The right upper lung lobe was collected, washed in PBS and maintained in 10% paraformaldehyde for 24 hours. Subsequently, histological sections of approximately 4-5 μm, which were stained with Hematoxylin-eosin (HE) or Masson’s trichrome (MT) for microscopic visualization and then photographed.

### CFU determination

The lung suspension was undergoing serial dilution in PBS and soon after it was seeded in petri dishes with Middlebrook 7H10 Agar (Becton Dickinson, US) supplemented with OADC enrichment medium (oleate, albumin, dextrose and catalase) (Difco, US). The plates were sealed and kept in an oven at 37º C and 5% CO2 for 21 days. After this period, colonies were counted to determine the CFU.

### Viral load determination

Viral loads in the experiments with the PR8 strain were defined by quantitative PCR (qPCR), comparing the amplification of lung supernatant samples with the amplification of viral cDNA from a sample of known concentration of PFU (standard curve method). The primers used for virus cDNA amplification were: PR8-NP, 5′ - GATTGGTGGAATTGGACGAT-3 ′ and 5′ -AGAGCACCATTCTCTCTATT-3′.

### Confocal microscopy

Paraffinized sections of approximately 10μm were made with the right upper lobe of the lung. After dewaxing, the sections were submitted to antigenic recovery. Subsequently, the sections blocked with 2% BSA in PBS for 30 min, and then incubated overnight at 4°C with primary anti-S100A9 antibody. After 24 hours, the sections were stained with fluorochrome-labeled anti-rabbit antibody and anti-CD4 and anti-IFN-γ conjugated antibodies. The slides were treated with the Sudan Black B (Sigma-Aldrich) reagent to remove the natural autofluorescence of the tissue.

Next, the slides were rinsed in PBS and coverslipped with ProLong Gold antifade reagent (Invitrogen). After being stained and treated, slides were washed, and coverslips were inserted. The high resolution SP5 confocal microscope (Leica) was used to view slides and image analysis performed in Fiji software.

### Mixed bone marrow chimera mice generation

C57BL/6 (CD45.2) mice were irradiated with 1000 rads and reconstituted with the 1:1 mixture of CD4-Cre (CD45.2) and CD4-Cre *P2rx7*^*fl/fl*^ (CD45.1/2) mouse bone marrow. Reconstitution was followed weekly by flow cytometry.

### RNA-sequencing analysis

WT or CD4-Cre *P2rx7*^*fl/fl*^ mice were infected with PR8/34. At day 7 post-infection, CD44^+^ CD4^+^ T cells from lungs (i.v.^+^ and i.v.^-^ compartments) and spleen were isolated by cell sorting. RNA was extracted using the RNeasy Plus Mini Kit (QIAgen). Library preparation and RNA-seq (DNBseq platform, PE 100bp pair-end read length) was done by BGI Americas. RNA-seq reads were mapped and raw count matrix was generated. DEG analysis was done using DESeq2, and genes with >2-fold changes and FDR <0.05 were considered for gene cluster analysis. Heatmaps, PCA plots and Pathway Enrichment Analyses were generated using the R-based BGI Dr.Tom online analysis toolkit (https://www.bgi.com/global/dr-tom/).

#### *In vivo* CXCR3 Inhibition

For anti-CXCR3 treatment, WT and *P2rx7*^*-/-*^ mice were infected with Influenza virus PR8/34 strain (1400 PFU) treated intraperitonially with 200μg of anti-mouse CXCR3 (BioXCell, Lebanon/NH, clone CXCR3-173) on days 0, 3, 4 and 6 p.i. On day 7 p.i. the lung was collected for analysis by flow cytometry.

#### *In vitro* experiments with human PBMC

For in vitro experiments using human samples the blood was collected by vacuum venipuncture in a tube containing anticoagulant heparin. PBMC were obtained using density gradient centrifugation with Ficoll and plated. The isolated PBMC from two individuals (AA and CC) previously characterized by having or not P2RX7 single-nucleotide polymorphisms (*rs3751143 A>C*) (Souza et al., 2021) were isolated and plated (1×10^5^). Both donors already had the active form of leprosy, caused by g/ Mycobacterium leprae. They were treated and are now healthy. To generate a specific response against mycobacteria, we activated the PBMC with mycobacterial extract (1 μg/mL) at the time of plating. Non-activated cells were used as a control. The cells were kept at 37°C and 5% CO_2_ and 24 h later part of the cells received supplementation (50 μM) of Adenosine 5-triphosphate (ATP - Sigma-Aldrich). After 24 h the cells were stained and subjected to flow cytometry analysis to assess the expression of Live/dead markers CD4, CD44, CXCR3 and Ki-67. Project submitted and approved to the Ethics and Research Committee of the Faculty of Medicine of Campos under CAAE number: 19679119.8.0000.5244.

### Quantification and statistical analysis

Details of the statistical analyses used for each experiment can be found in the figure legends. Statistical analyses were performed using the GraphPad Prism 9 software. Data were described as mean with error bars indicating the SD T-tests were used to assess differences between only two groups. One-way ANOVA tests and Tukey’s post hoc tests were used to assess the effects of only 1 parameter among more than 2 groups. Differences between groups were considered significant when p < 0.05 (*), p < 0.01 (**), p < 0.001 (***) or p < 0.0001 (****).

## SUPPLEMENTAL INFORMATION

**Figure S1.**
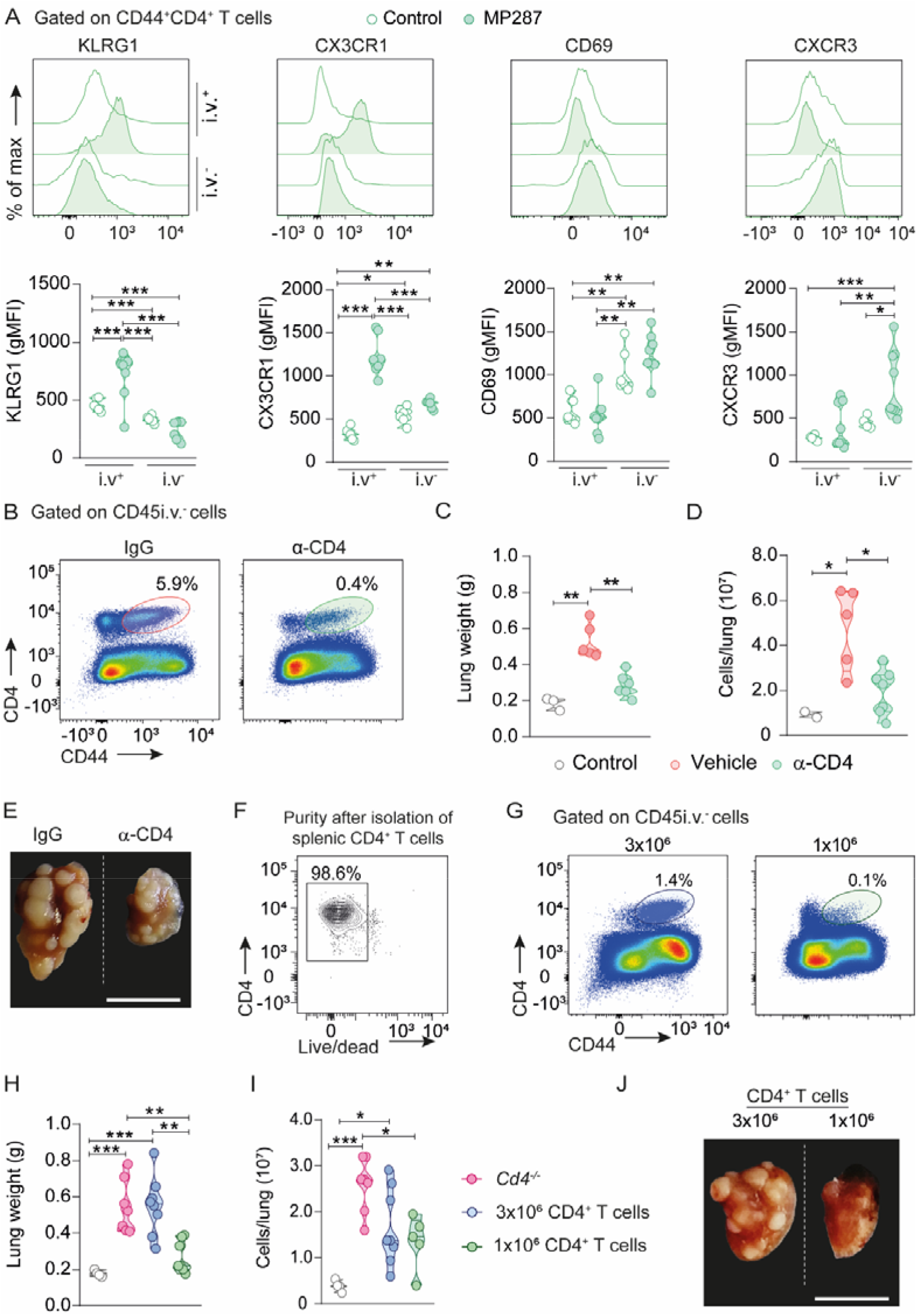
Effects of P2RX7 on lung CD4^+^ T cells and of CD4^+^ T cells on lung TB pathology caused by hypervirulent mycobacteria. (**A-J**) C57BL/6 (WT) and CD4-KO (*Cd4*^*-/-*^) mice were infected with ∼100 CFU of the *Mbv*-MP287 strain. Non-infected mice were used as controls. At 28 days p.i., mice were injected i.v. with anti-CD45 fluorescent antibodies 3 minutes before lung harvest. (**A**) Representative flow cytometry histograms (top panels) and violin plots (bottom panels, gMFI average) of the expression of KLRG1, CXC3CR1, CD69 and CXCR3 on CD45i.v.^+^ and CD45i.v.^-^ lung CD4^+^ T cells of WT mice. (**B**) Pseudocolor plots of CD4 and CD44 expression of on lung CD45i.v.^-^ cells of WT mice. (**C**) Lung weight values of WT mice. (**D**) Total cells per lung of WT mice. (**E**) Macroscopic images of right upper lung lobes of WT mice (bar scales = 1 cm). (**F**) Contour plots showing the purity and viability (Live/Dead) of splenic CD4^+^ T cells. (**G**) Pseudocolor plots of CD4 and CD44 expression on lung CD45i.v.^-^ cells of infected *Cd4*^-/-^ mice transferred with CD4^+^ T cells. (**H**) Lung weight values of mice described in G. (**I**) Total cells per lung of mice described in G. (**J**) Macroscopic images of right upper lung lobes of mice described in G. Data are from 2–3 independent experiments; n = 3-5 per experimental group. Graphical data is shown as means ± SD. p values from One-way ANOVA are indicated by ^*^, p <0.05; ^**^, p<0.01; ^***^, p <0.001.

**Figure S2.**
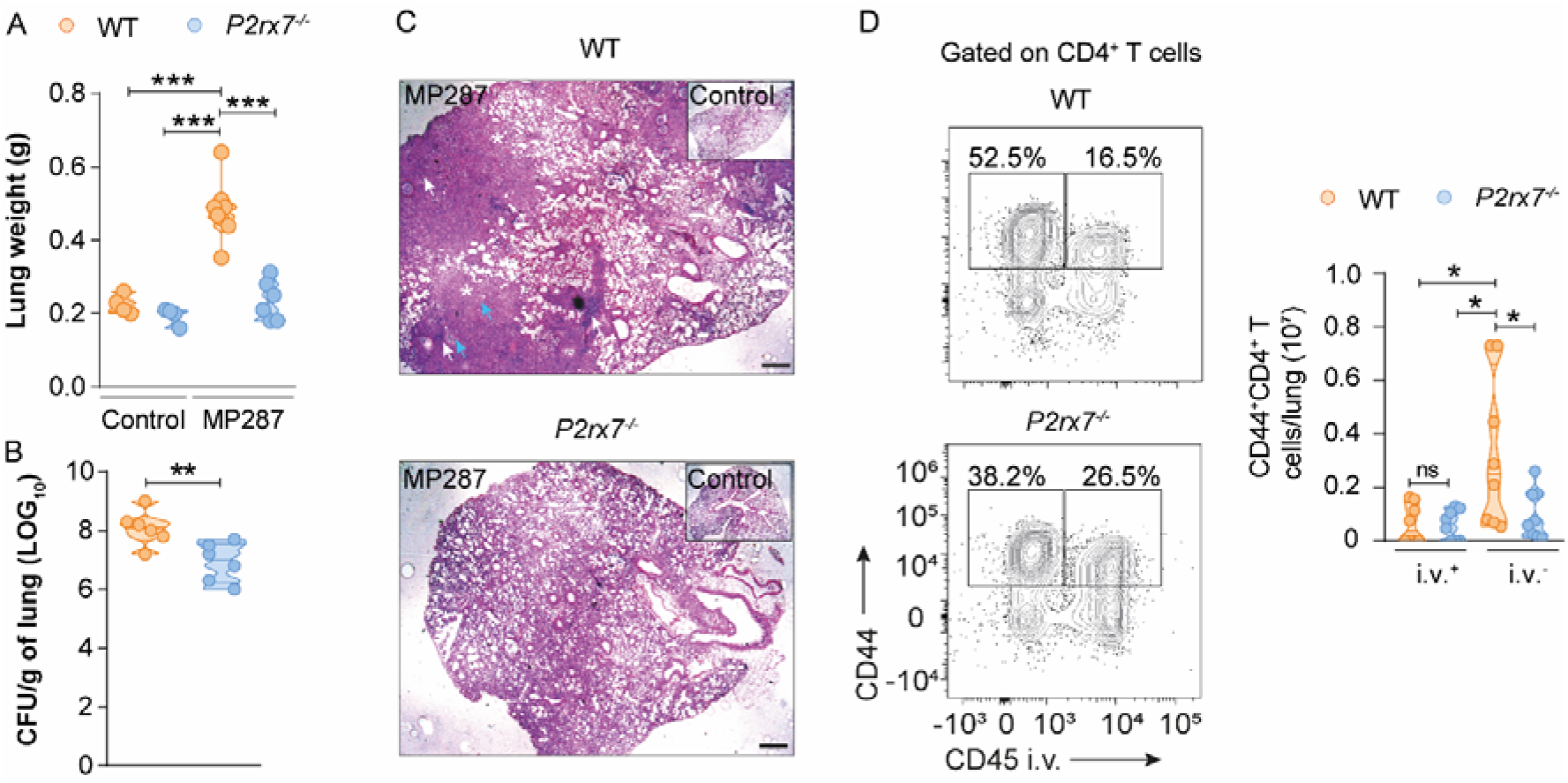
Effects of P2RX7 on TB caused by hypervirulent mycobacteria. WT and P2RX7-KO (*P2rx7*^-/-^) mice were infected with ∼100 CFU of the *Mbv*-MP287 strain. Non-infected mice were used as controls. At 28 days p.i., mice were injected i.v. with anti-CD45 fluorescent antibodies 3 minutes before lung harvest. (**A**) Lung weights. (**B**) CFU numbers per lung. (**C**) Representative lung sections stained with HE (bar scales = 500 μM). White asterisks indicate areas of necrosis, white arrows indicate area of alveolitis and blue arrows indicate bronchial obstruction. (**D**) Left: representative flow cytometry plots of CD44 and CD45i.v. expression in CD4^+^ T cells. Right: violin plots of the number of CD45iv^+^ and CD45iv^-^ CD44^+^CD4^+^ T cells per lung. Data are from 2 independent experiments; n = 3-5 per experimental group per experiment. Graphical data is shown as means ± SD. p values from One-way ANOVA (Tukey post-tests) (**A** and **D**) or unpaired T-tests (**B**) are indicated by ^*^, p <0.05; ^**^, p<0.01; ^***^, p <0.001.

**Figure S3.**
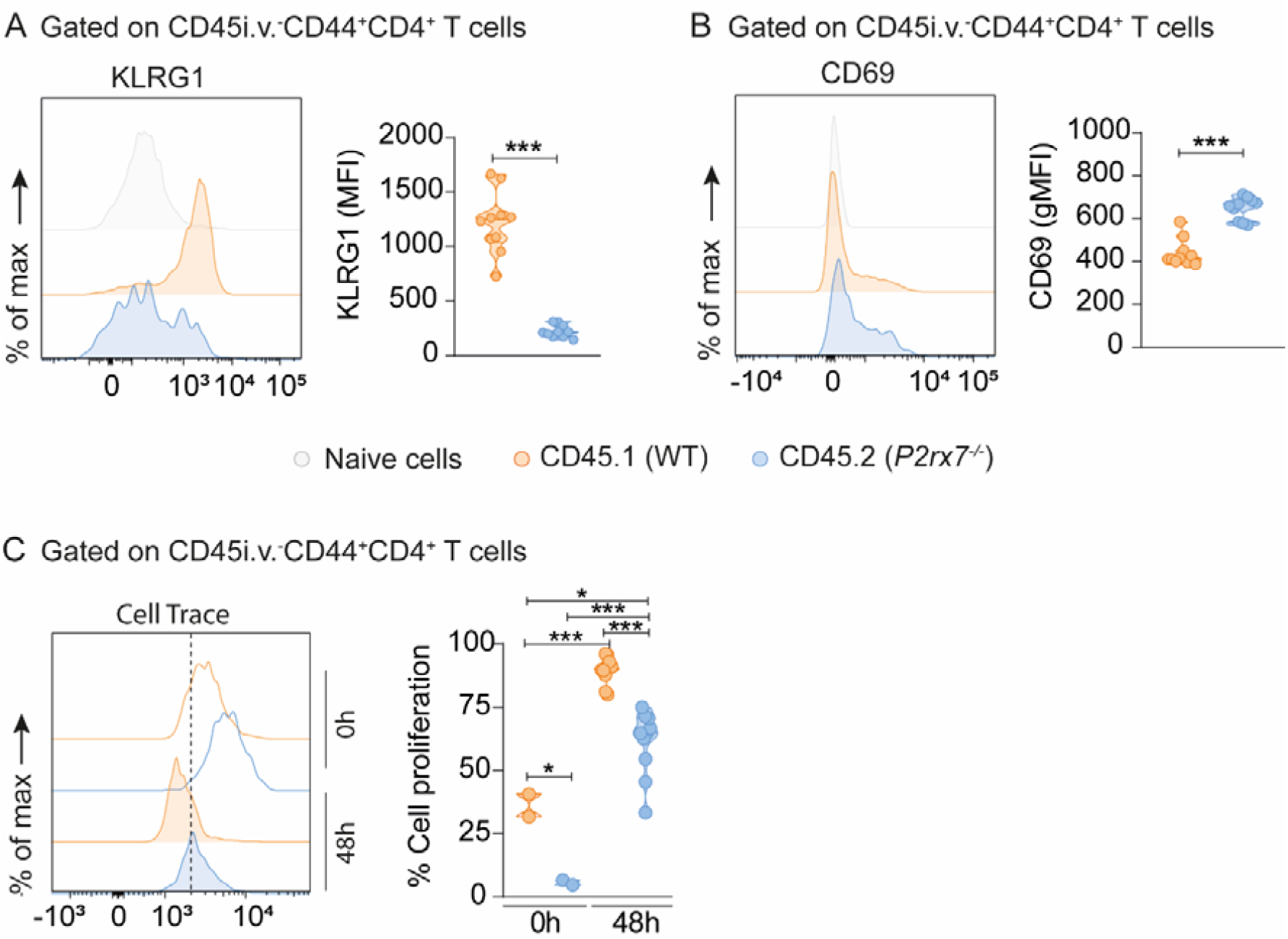
Effects of P2RX7 in the phenotype and proliferation of parenchymal and intravascular CD4^+^ T cells in response to MP287 infection. CD4-KO (*Cd4*^*-/-*^) mice were infected with ∼100 CFU of the *Mbv*-MP287 strain. On day 5 p.i. splenic CD4^+^ T cells (1×10^6^) from WT (CD45.1^+^) and P2RX7-KO (CD45.2^+^; *P2rx7*^*-/-*^) mice were co-transferred (1:1) to infected *Cd4*^*-/-*^ mice. At 28 days p.i., mice were injected i.v. with anti-CD45 fluorescent antibodies 3 minutes before lung harvest. (**A**) Representative histogram and average Geometric Mean Fluorescence Intensity (gMFI) values of KLRG1 xpression in CD45i.v.^-^ CD44^+^CD4^+^ T cells and naive CD4^+^ T cells. (**B**) Representative histograms and Geometric Mean Fluorescence Intensity (gMFI) values of CD69 expression in CD45i.v.^-^CD44^+^CD4^+^ T cells and naive CD4^+^ T cells. (**C**) *Ex vivo* cell tracer in CD45i.v.^-^CD44^+^CD4^+^ T cells 0 and 48 h after staining. Data are from 2 independent experiments; n =5 per experimental group per experiment. Graphical data is shown as means ± SD p values from One-way ANOVA (Tukey post-tests) (**A and B**) or unpaired T-tests (**C**) are indicated by ^*^, p <0.05; ^**^, p<0.01; ^***^, p <0.001.

**Figure S4.**
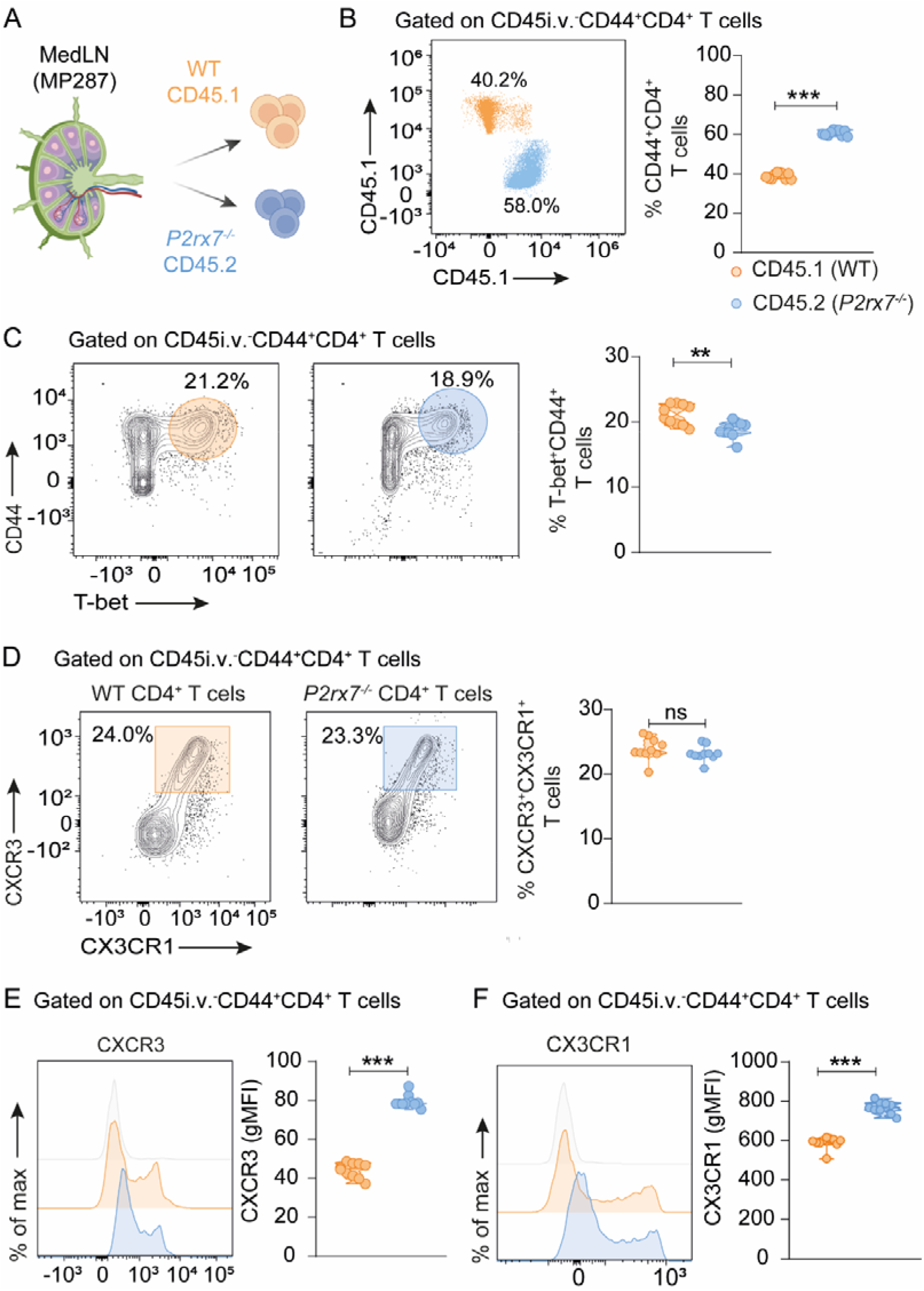
Cell-intrinsic P2RX7 is not required for CD4^+^ T cell activation and effector differentiation in lung-draining lymph nodes in response to severe TB. CD4-KO (*Cd4*^*-/-*^) mice were infected with ∼100 CFU of the *Mbv*-MP287 strain. On day 5 p.i. splenic CD4^+^ T cells (1×10^6^) from WT (CD45.1^+^) and P2RX7-KO (CD45.2^+^; *P2rx7*^*-/-*^) mice were co-transferred (1:1) to infected *Cd4*^*-/-*^ mice. At 28 days p.i., mice were injected i.v. with anti-CD45 fluorescent antibodies 3 minutes before lung and mediastinal lymph nodes (medLNs) harvest. (**A**) Schematic illustration of the experimental protocol (Created with BioRender.com). (**B**) Left: representative flow cytometry plots of CD45.1 and CD45.2 expression on CD45i.v.^-^CD44^+^CD4^+^ T cells. Right: frequencies (%) of CD45i.v.^-^ CD44^+^CD4^+^ T cells in medLNs. (**C**) Left: representative flow cytometry plots of CD44 and T-bet expression on CD45i.v.^-^CD4^+^ T cells in medLNs. Right: frequencies (%) of T-bet^+^ cells. (**D**) Left: representative flow cytometry plots of CXCR3 and CX3CR1 expression on CD45i.v.^-^CD44^+^CD4^+^ T cells in medLNs. Right: frequencies (%) of CXCR3^+^ and CX3CR1^+^ cells. (**E**) Representative flow cytometry plots and gMFI values of CXCR3 and CX3CR1 expression on CD45i.v.^-^CD44^+^CD4^+^ T cells in medLNs. Data are from 3 independent experiments; n =5 per experimental group per experiment. Graphical data is shown as means ± SD. p values from Unpaired T-test are indicated by ^*^, p <0.05; ^**^, p<0.01; ^***^, p <0.001.

**Figure S5.**
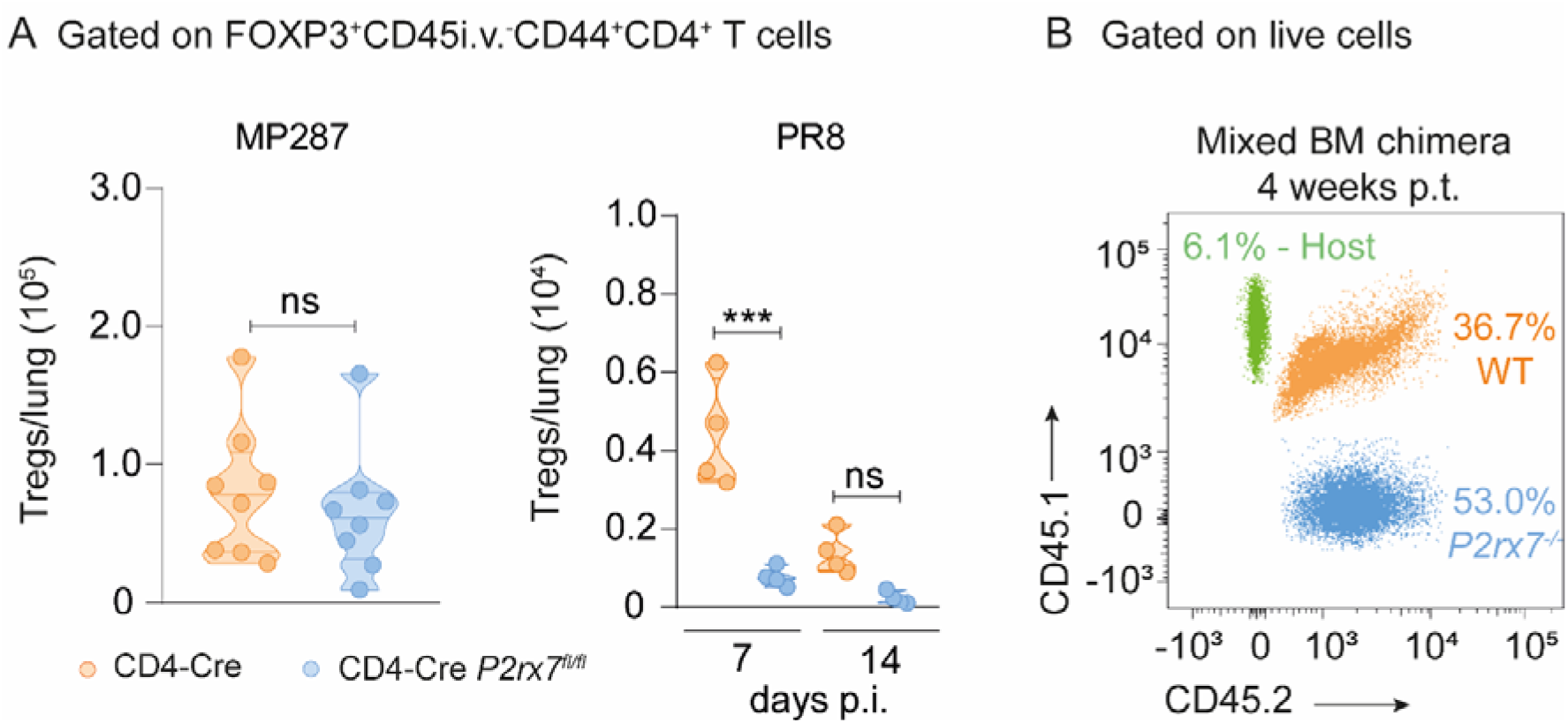
Effects of P2RX7 expression on Treg number during MP287 and PR8 infection and reconstitution of mixed bone marrow chimera mice. WT (CD4-Cre) or T cell-P2RX7-KO (CD4-Cre *P2rx7*^*fl/fl*^) mice were infected with MP287 (∼ 100 CFU) or PR8 (∼1400 PFU) strains. At 28 days p.i., mice were injected i.v. with anti-CD45 fluorescent antibodies 3 minutes before tissue harvest. (**A**) Left: the violin plots show the number of FOXP3^+^CD45i.v.^-^CD44^+^CD4^+^ T cells per MP287-infected lung. Right: the violin plots show the number of FOXP3^+^CD45i.v.^-^CD44^+^CD4^+^ T cells per PR8-infected lung. (**B**) Representative flow cytometry plots showing CD45.1 and CD45.2 expression on live cells of mixed BM chimera mice blood 4 weeks post transfer. Data are from 2 independent experiments; n = 2-4 per experimental group per experiment. Graphical data is shown as means ± SD. p values from unpaired T-tests are indicated by ^*^, p <0.05; ^**^, p<0.01; ^***^, p <0.001.

**Figure S6.**
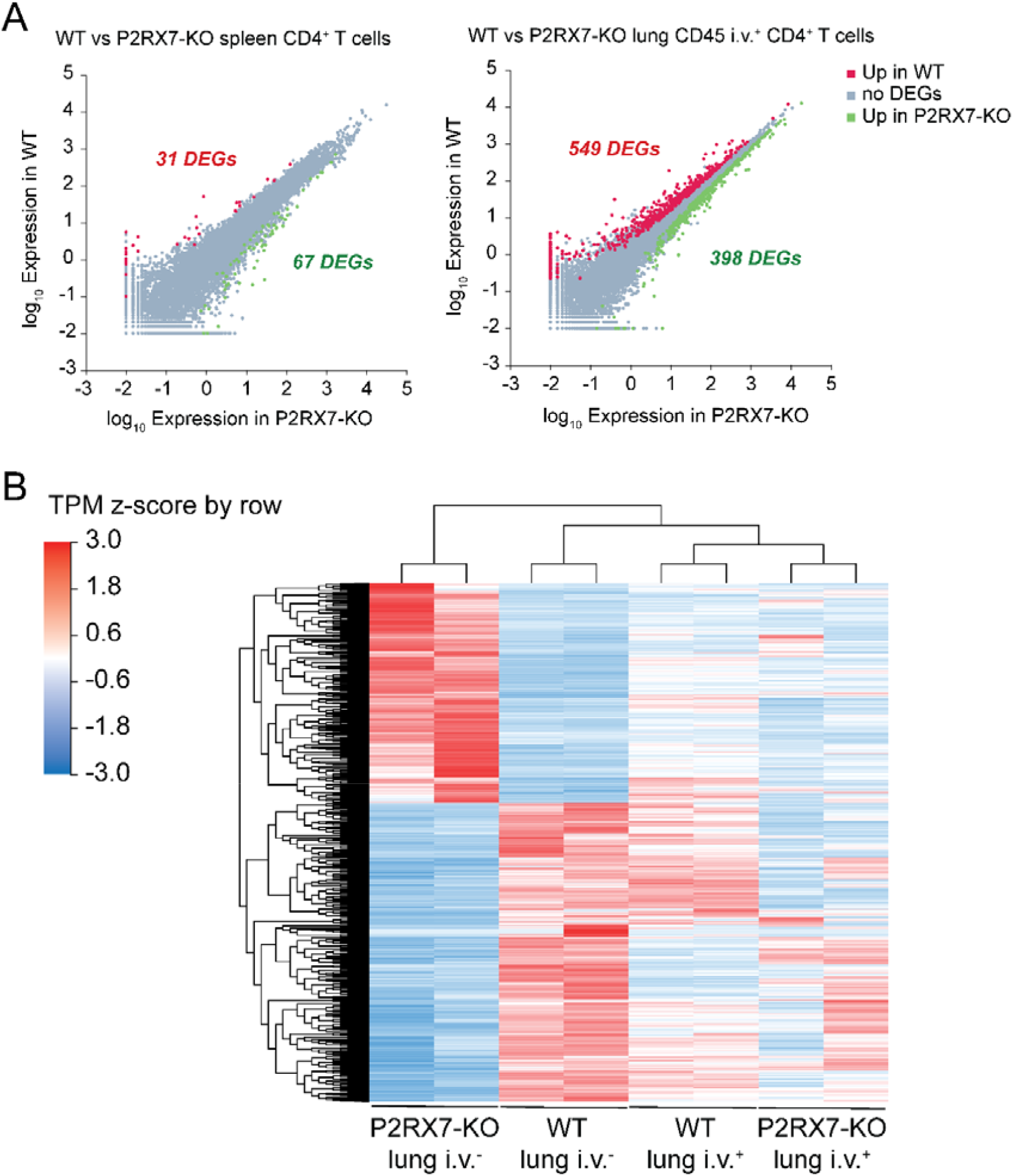
Transcriptional signatures of WT and P2RX7-KO lung and spleen CD4^+^ T cells responding to acute influenza infection. (**A-B**) WT (CD4-Cre) and T cell-P2RX7-KO (CD4-Cre *P2rx7*^fl/fl^) mice were infected with influenza (PR8, 1400 PFU). At day 7 post-infection, lung i.v.^-^, lung i.v.^+^ and spleen CD44^+^ CD4^+^ T cells were sorted and RNA-seq analysis was performed. (**A**) Scatter plots with DEGs highlighted between WT and P2RX7-KO spleen (left) and lung i.v.^+^ (right) CD4^+^ T cells. (**B**) Heatmap displaying, for all lung subsets analyzed, the DEGs between WT and P2RX7-KO lung i.v.^-^ CD4^+^ T cells. Each replicate is a pool of sorted CD4^+^ T cells from five mice; n=2 replicates per experimental group.

